# Proteoforms of the SARS-CoV-2 nucleocapsid protein are primed to proliferate the virus and attenuate the antibody response

**DOI:** 10.1101/2020.10.06.328112

**Authors:** Corinne A. Lutomski, Tarick J. El-Baba, Jani R. Bolla, Carol V. Robinson

## Abstract

The SARS-CoV-2 nucleocapsid (N) protein is the most immunogenic of the structural proteins and plays essential roles in several stages of the virus lifecycle. It is comprised of two major structural domains: the RNA binding domain, which interacts with viral and host RNA, and the oligomerization domain which assembles to form the viral core. Here, we investigate the assembly state and RNA binding properties of the full-length nucleocapsid protein using native mass spectrometry. We find that dimers, and not monomers, of full-length N protein bind RNA, implying that dimers are the functional unit of ribonucleoprotein assembly. In addition, we find that N protein binds RNA with a preference for GGG motifs which are known to form short stem loop structures. Unexpectedly, we found that N undergoes proteolytic processing within the linker region, separating the two major domains. This process results in the formation of at least five proteoforms that we sequenced using electron transfer dissociation, higher-energy collision induced dissociation and corroborated by peptide mapping. The cleavage sites identified are in highly conserved regions leading us to consider the potential roles of the resulting proteoforms. We found that monomers of N-terminal proteoforms bind RNA with the same preference for GGG motifs and that the oligomeric state of a C-terminal proteoform (N_156-419_) is sensitive to pH. We then tested interactions of the proteoforms with the immunophilin cyclophilin A, a key component in coronavirus replication. We found that N_1-209_ and N_1-273_ bind directly to cyclophilin A, an interaction that is abolished by the approved immunosuppressant drug cyclosporin A. In addition, we found the C-terminal proteoform N_156-419_ generated the highest antibody response in convalescent plasma from patients >6 months from initial COVID-19 diagnosis when compared to the other proteoforms. Overall, the different interactions of N proteoforms with RNA, cyclophilin A, and human antibodies have implications for viral proliferation and vaccine development.

## Introduction

Severe acute respiratory syndrome coronavirus 2 (SARS-CoV-2) is the etiological agent of coronavirus disease 2019 (COVID-19) which reached pandemic status in fewer than three months following its discovery. As of October 2020, there are >34 million infected and more than 1 million deaths.^1^ However, SARS-CoV-2 is not the first coronavirus to cause widespread disease. Two epidemics in the last two decades have been caused by coronaviruses: severe acute respiratory syndrome (SARS) in 2002 and the Middle East respiratory syndrome (MERS) in 2012. Despite belonging to the same viral genus, SARS-CoV-2 has proven far more infectious, implying molecular differences between the viruses. Understanding the molecular components of viral proliferation in SARS-CoV-2 is therefore important to develop new therapeutics to combat COVID-19.

SARS-CoV-2 packages a large RNA genome of ~30 kb which encodes for 25 non-structural and four structural proteins (spike, nucleocapsid, membrane, and envelope proteins). The nucleocapsid (N) protein is one of the most abundant viral proteins; hundreds of copies of N make up the viral core that encapsulates the genomic RNA. In addition to its essential role in RNA replication/transcription and virion assembly, coronavirus N proteins play essential roles in host cell signaling pathways, immune system interference, cell cycle regulation, and chaperone activity.^2^ Interestingly, the nucleocapsid and spike proteins are the main immunogens circulating in the blood of COVID-19 patients.^3^ However, quantitative measurements of plasma or serum from SARS-CoV-2 patients found that the adaptive immune response to the N protein is more sensitive than the spike protein^4^, making it a better indicator of early disease and a good target for antiviral therapies.

N protein has been the subject of much effort in structure elucidation in order to guide the design of novel antivirals.^5,6^ Several compounds targeting nucleocapsid proteins of other viruses have proven effective in *in vitro* studies, specifically blocking replication, transcription, and assembly.^7,8,9,10^ However, the timeline for developing approved antivirals takes years and the need for treatments to combat COVID-19 is urgent. Therefore, repurposing existing drugs is an attractive and immediate solution to combat COVID-19.^11^ On this front, key host interactions have been mapped for the proteins encoded by SARS-CoV-2 to identify host targets for drug repurposing.^12^ One particular study revealed 66 potential drug targets, three of which are direct interactors of N protein. The safe and effective use of new and existing therapeutics to target specific viral processes however relies on an understanding of the sequence of interactions between viral and host proteins and their controls.

Here, we present a comprehensive analysis of the SARS-CoV-2 N protein using native mass spectrometry (MS), top-down fragmentation, and bottom-up sequencing. We find that the full-length N protein undergoes proteolysis at highly conserved sites to generate at least five unique proteoforms. We identify various stoichiometries of the proteoforms that are influenced by pH and that may be critical targets in drug design. We evaluate the propensity for N and N proteoforms to bind different RNA sequences and find that only the dimeric form of full-length N binds to RNA, suggesting it is the functional unit of ribonucleoprotein assembly. In addition, we show that two N proteoforms directly interact with cyclophilin A, a highly abundant cytosolic host protein implicated in viral replication. We found that cyclosporin A, an immunosuppressive drug, abolishes the interaction between N proteoforms and cyclophilin A. Finally, using convalescent plasma from patients >6 months from initial COVID-19 diagnosis, we found that the antibody response to the N-terminal proteoforms was significantly attenuated compared to the full-length N protein. Interestingly, the antibody response for full-length protein and the C-terminal proteoform were not statistically different, suggesting that the antigenic site for antibody recognition is localized to the C-terminus of the N protein. Our results indicate that SARS-CoV-2 N protein is a multifunctional protein and propose that N proteoforms are primed for additional functions related to viral propagation.

## Results and Discussion

### N Protein undergoes proteolysis in the vicinity of the linker region

Nucleocapsid proteins of coronaviruses share a similar topological organization^13^ and show high sequence homology among related coronaviruses.^14^ The N protein is characterized by two major structural domains, the RNA binding and oligomerization domain (Figure 1A). ^15^ Similar to other coronavirus nucleocapsid proteins, the two domains are separated by a long and flexible linker region thought to be devoid of secondary structure.^2^ An N-terminal arm and a C-terminal tail flank the RNA binding and oligomerization domains, respectively. We constructed a plasmid consisting of the full-length nucleocapsid protein, an N-terminal purification tag, and a cleavage site (Figure 1A). We expressed and purified this construct in *Escherichia coli* and verified the cleavage of the affinity tag via SDS-PAGE (Figure 1B). Before removal of the purification tag, three distinct protein bands were detected at ~49, ~38 and ~28 kDa. Following removal of the tag, all three protein bands migrated by ~3 kDa, or the mass of the tag (Table 1).

**Table 1.** Deconvoluted and sequence masses of N_FL_ and N proteoforms.

| **Protein** | **Deconvoluted Mass<sup>a</sup> ± s.d. (Da)** | **Sequence Mass (Da)** |
| --- | --- | --- |
| N-terminal tagged N<sub>FL</sub> | -- | 48,859.21 |
| N<sub>FL</sub> | 45,769 ± 2 | 45,769.83 |
| N<sub>FL</sub> dimer | 91,537 ± 2 | 91,539.66 |
| N<sub>156-419</sub> | 28,735 ± 2 | 28,696.12 |
| N<sub>156-419</sub> dimer | 57,534 ± 1 | 57,392.24 |
| N<sub>1-209</sub> | 22,611 ± 1 | 22,612.71 |
| N<sub>1-220</sub> | 23,540 ± 0.3 | 23,541.73 |
| N<sub>1-273</sub> | 29,402 ± 0.6 | 29,382.43 |
| N<sub>1-392</sub> | 42,922 ± 1 | 42,918.74 |

<sup>a</sup> Determined using at least three adjacent charge states

**Figure 1.**
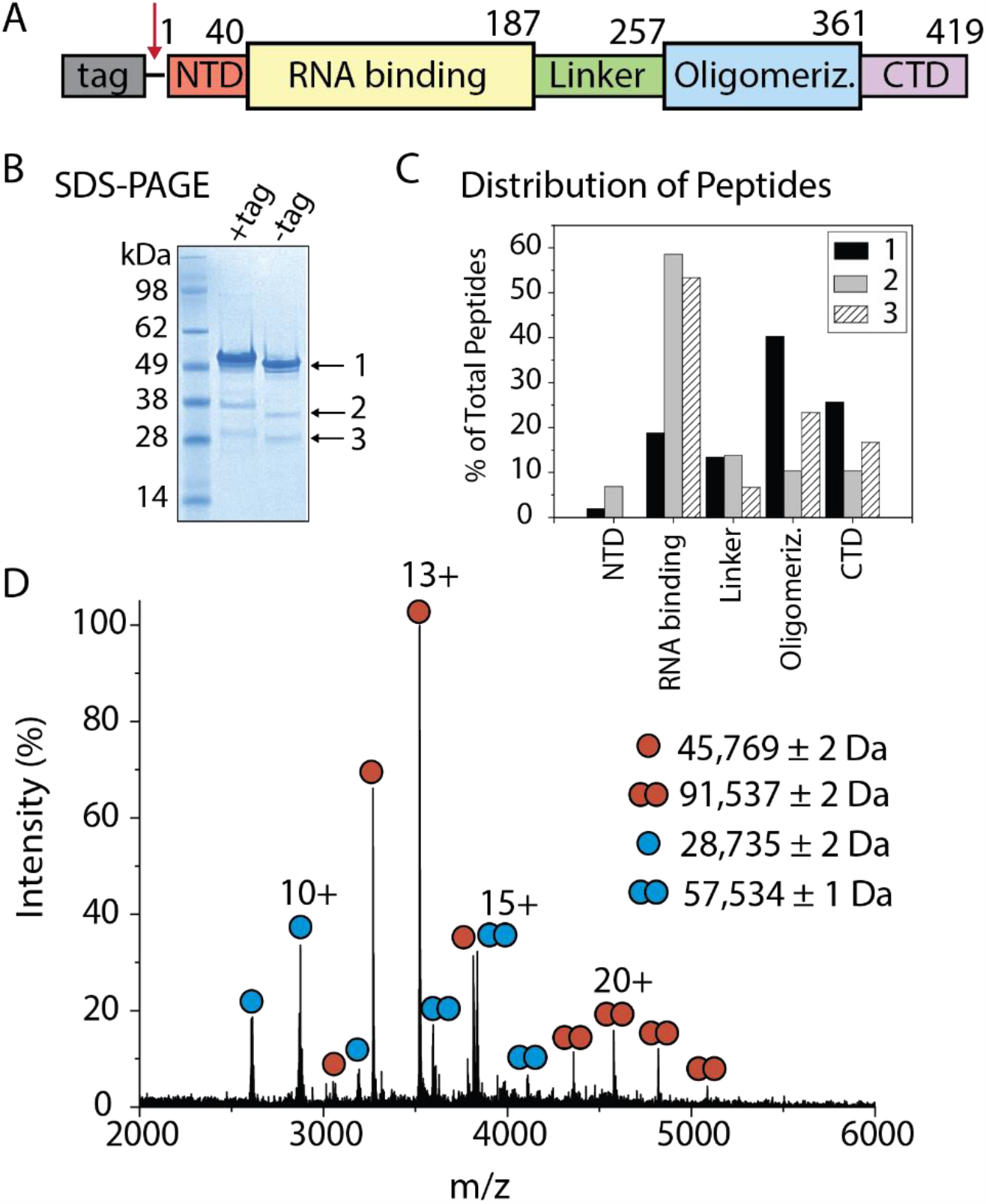
SARS-CoV-2 N protein exists as an ensemble of proteoforms. (A) Scheme depicting the full-length construct for expression in *E. coli*. The construct contains an N-terminal purification tag that is cleaved at the site indicated (red arrow). (B) SDS-PAGE of the N protein construct in (A) before and after tag cleavage. Three distinct protein bands (denoted 1-3) are observed in lanes labeled +tag and −tag. (C) The protein bands were subjected to in-gel digestion with trypsin followed by LC-MS based proteomics. (C) Histogram displaying percentage of peptides detected, relative to the total peptide count, across the five protein domains: N-terminal domain (NTD), RNA binding, Linker, Oligomerization, and C-terminal Domain (CTD). (D) Mass spectrum of the N protein after enrichment by size exclusion chromatography shows that, despite extensive purification, N protein exists as an ensemble of proteoforms. Four charge state distributions correspond to monomers and dimers of full-length N protein (red circles, MW 45,769 Da) and a proteoform of N protein (blue circles, MW 28,735 Da).

We confirmed that each band corresponded to N protein by in-gel trypsin digestion followed by LC-MS-based bottom-up proteomics. All three bands contained peptides from the N protein, resulting in 78.1%, 49.9%, and 43.2% sequence coverage for bands 1, 2, and 3, respectively. To determine the representation of protein domains in each gel band, we plotted the distribution of the peptides detected across the five protein domains (Figure 1C). As anticipated, we observe an unbiased distribution of peptides across all five domains for band 1, consistent with the expectation that tryptic peptides would be reasonably distributed across the full-length protein. Over 50% of the total peptides detected in bands 2 and 3 were localized to the RNA binding domain, suggesting that the proteins are predominantly N-terminal derivatives. However, in all three bands, peptides located in the 58-residue C-terminal domain were detected, indicating the purified protein is made up of a diverse mixture of N proteoforms^16^ and not biased to N-terminal species due to the location of the purification tag.

We recorded a native mass spectrum to identify the masses of the proteoforms present together with the full-length protein (Figure 1D). Two main charge state envelopes centered at 13+ and 20+ species were identified, consistent with coexistence of monomers and dimers. Deconvolution of the *m/z* signals provided experimental masses of 45,769 ± 1 Da and 91,537 ± 2 Da, respectively, which are in excellent agreement with the theoretical monomeric (45,769.83 Da) and dimeric (91,539.66 Da) masses of full-length N protein. The second distribution of monomers and dimers have deconvoluted masses of 28,735 ± 2 Da and 57,534 ± 1 Da, respectively. The SDS-PAGE analysis indicated the presence of additional proteoforms not visible in this spectrum. We separated these lower molecular weight species from the full-length N using size exclusion chromatography (Figure S1). The mass spectrum of the pooled fractions revealed the presence of four additional proteoforms that range in mass from 22,612 to 42,922 Da.

Over several days, the mass spectrum evolved to reveal a series of peaks corresponding to five unique protein distributions (Figure 2A). To determine the identity of each proteoform, we adapted a two-tiered tandem mass spectrometry approach^17^ to determine intact mass and amino acid sequence for each series of peaks in the mixture. An individual peak in the mass spectrum was first isolated and subjected to electron transfer dissociation (ETD) under conditions that do not result in the formation of fragments but instead produce a series of charge-reduced peaks (Figure 2B). The charge-reduced spectrum was necessary to confirm the assignment of the charge state series for each proteoform. The assigned charge states were then used to obtain deconvoluted masses of each proteoform present in solution.

**Figure 2.**
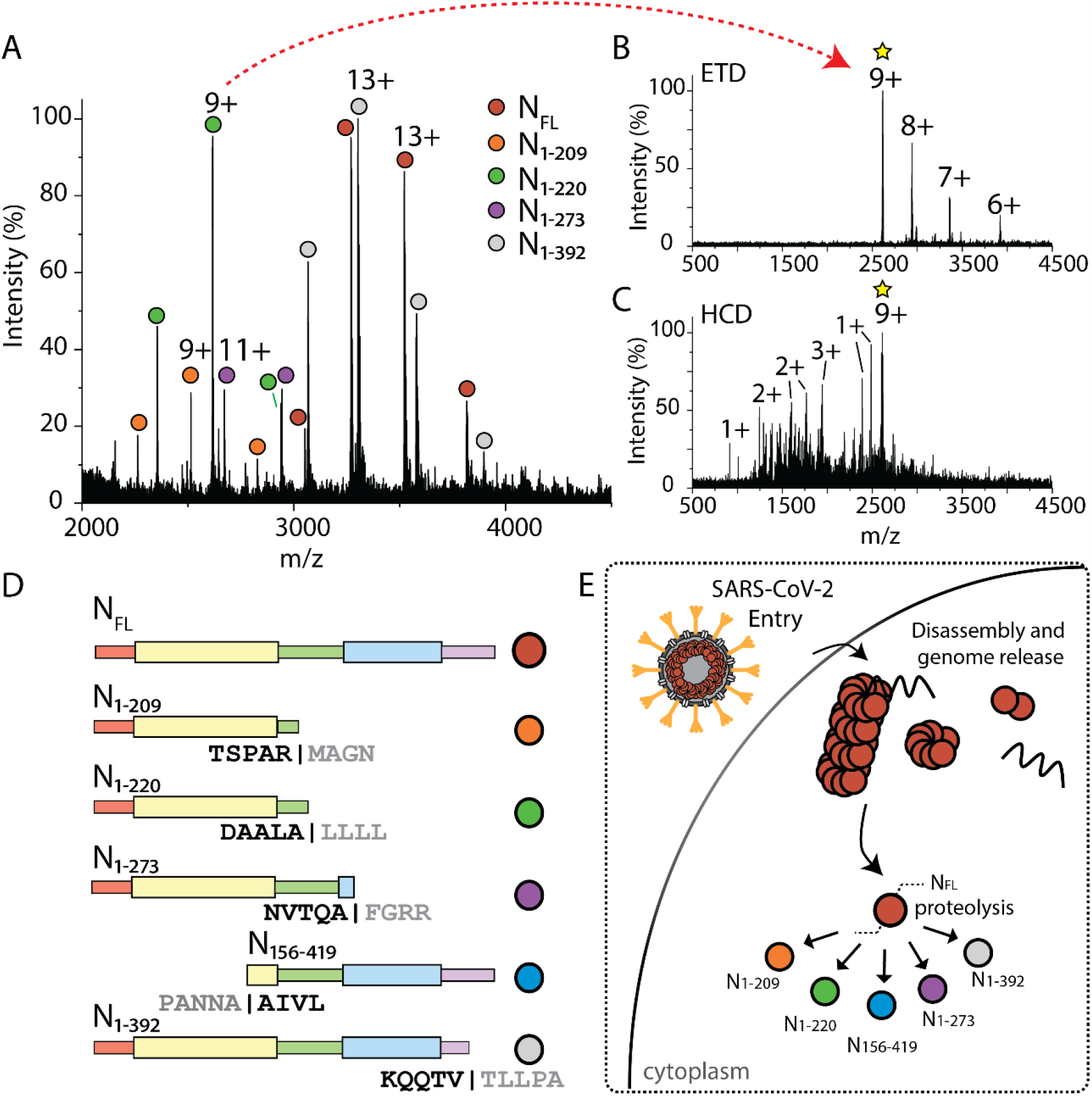
N protein undergoes proteolysis in a highly specific manner. (A) Mass spectrum of N protein after incubation with protease inhibitors for one week reveals the coexistence of five distinct charge state distributions corresponding to N proteoforms. The chemical composition of each proteoform was determined using top-down MS (B,C). (B) Charge-reduced mass spectrum resulting from electron-transfer dissociation (ETD) of the selected 9+ charge state at *m/z* 2616.79. (C) Mass spectrum of sequence ions for the same parent ion generated by higher-energy collision induced dissociation (HCD). (D) Scheme representing the composition of protein domains for the observed proteoforms as determined by top-down MS. Five distinct proteoforms are observed: N_1-209_, N_1-220_, N_1-273_, N_156-419,_ and N_1-392_. The exact site of cleavage, including the five residues flanking either side of each cleavage site, is indicated below each construct. (E) Scheme depicting the proteolytic cleavage of full-length N protein after virus entry and nucleocapsid disassembly.

We tentatively assigned the proteoforms based on their intact masses and then generated sequence ions to confirm our assignment. Individual proteoforms were subjected to fragmentation by higher-energy collision induced dissociation (HCD), which accelerates the isolated ions into an inert gas to induce fragmentation along the amide backbone. The fragmentation products were then used to determine the molecular composition and to localize the exact site of cleavage. The HCD spectrum results in a series of singly-, doubly-, and triply-charged sequence ions exemplified by the fragmentation of the 9+ charge state at *m/z* 2616.79, Figure 2C. Fragmentation of intact proteins under native conditions is expected to yield 3-10% sequence coverage at the termini^18^ and the propensity for fragmentation differs depending on several criteria (e.g. mass, charge, composition, structure) for natively folded proteins.^19^ Here, we achieved ~4% sequence coverage of the proteoforms by native top-down MS (Table S1). Fragmentation at the termini was complementary to our goal – to confirm sites of cleavage that result in distinct proteoforms of N protein.

### Cleavage sites are highly conserved in the SARS-CoV-2 genome

Considering the proteoforms identified, we note that three result from cleavage after alanine (residue pairs A|L, A|F, and A|A), one from cleavage following an arginine (R|M), and one results from cleavage in the C-terminal tail following a valine (V|T), (Figure 2D). Proteoforms N_1-209_ and N_1-220_ contain primarily the RNA binding domain as they cleave within the flexible linker region. The major component of N_1-273_ is also the RNA binding domain, but in this case is followed by the linker region and a small portion of the oligomerization domain. Conversely, N_156-419_ comprises mainly the oligomerization domain while still retaining a small portion of the RNA binding domain.

Although we find no sequence similarity in the residues that flank each cleavage site, a commonality is that cleavage occurs immediately adjacent to a hydrophobic residue. We were intrigued to discover if the cleavage sites were subject to mutation. The locations of the gene and amino acid mutations have been mapped for 38,318 SARS-CoV-2 genome sequences obtained from the China National Center for Bioinformation 2019 Novel Coronavirus Resource.^15^ In the sampled population, there were 200 amino acid mutations that occurred more than once, and these substitutions were heavily localized at or near the linker region. The proteolytic sites at A220, A273, A156, T392 were mutated only 4, 1, 9, and 2, times, in >38,000 genomes sequenced, making them highly conserved. By comparison, the site at R209 was mutated a total of 36 times; most commonly mutated to T via a G→C missense mutation at position 626 in the gene. The specificity of cleavage at the conserved residues identified here, despite no common motif, suggest that there may be a structural component directing proteolysis, however the exact mechanism is unclear. This is supported by small angle x-ray scattering measurements which demonstrate that the flexible linker region is not fully extended but contains elements of structure.^20^

### The oligomeric states of N proteoforms are influenced by pH

To better understand the role of the individual proteoforms we expressed and purified four individual constructs and recorded native mass spectra of N_1-209_, N_1-220_, N_1-273_, and N_156-419_ from solutions at different pH (Figures S3-S6). Mass spectra for N_1-209_ and N_1-220_ recorded at pH 5.0, 7.4 and 8.0 reveal highly abundant charge state series centered at 9+ and a low abundance distribution of signals centered at the 13+ charge state corresponding to monomers and dimers, respectively. The mass spectra for N_1-273_ show that it is predominantly monomeric with no significant change in charge state distribution or oligomeric state across the range of pH values tested. However, we observe two low abundant charge state series corresponding to N_1-209_ and N_1-220_, suggesting that N_1-273_ continues to undergo cleavage at the previously mapped residues (Figure S6).

In contrast to the N-terminal proteoforms, mass spectra for N_156-419_ reveal multiple charge state distributions at pH 5.0, 7.4 and 8.0 (Figure S6). At pH 8, the mass spectrum reveals three charge state distributions centered at 10+, 18+, and 27+ with average masses corresponding to monomers, trimers, and a low population of hexamers (Table S2). At pH 7.4, monomers, dimers, and trimers persist. At the lowest pH (pH 5.0) N_156-419_ is exclusively trimeric. Finally, the mass spectra for full-length N at pH 5.0 and 8.0 reveal broadened and featureless peaks suggesting that N_FL_ is likely aggregated (Figure S7). A low abundance series of highly charged peaks centered at 18+ at pH 8.0 indicates some protein unfolding. Overall, we conclude that N_1-209_, N_1-220,_ N_1-273_ do not undergo significant pH-dependent changes in oligomeric state while C-terminal fragments are highly sensitive with trimers predominating under both high and low pH conditions.

### RNA sequence influences binding stoichiometry

RNA binding and ribonucleoprotein complex formation is the primary function of coronavirus N proteins. The N protein binds nucleic acid nonspecifically^20^, however the production of infectious virions relies on N protein forming specific interactions with viral RNA among an abundance of different cellular RNA species. With knowledge of the stoichiometries of N_FL_ and N proteoforms, we sought to determine the propensity for these species to bind specific RNA sequences. We created single-stranded RNA oligonucleotides comprised of 20 nucleotides of repeating sequences (4×-GAUGG, 4×-GAGAA). Considering the promiscuity of N protein, we chose sequences shown to interact with the human immunodeficiency virus (HIV) polyprotein (which includes a nucleocapsid domain) at the different stages of virus assembly^21^ and hypothesized that N proteins of different oligomeric states would exhibit similar bias toward artificial RNA motifs.

We incubated N proteoforms and RNA oligonucleotides at a molar ratio of 4:1 protein:RNA and recorded native mass spectra for all N protein-RNA complexes. The mass spectrum for N_1-209_ bound to 4×-GAUGG (Figure 3A) reveals three charge state distributions with masses that correspond to apo N_1-209_ monomer and N_1-209_ bound to one and two 4×-GAUGG RNA oligonucleotides. The charge state distribution corresponding to N_1-209_ bound to two 4×-GAUGG RNA predominates over the single RNA bound protein. Conducting the same experiment with a different oligonucleotide (4×-GAGAA) reveals only one additional charge state distribution for N_1-209_ bound to one oligonucleotide (Figure 3B). Similar RNA binding stoichiometries are observed for N_1-220_ and N_1-273_; the mass spectra for N_1-220_ and N_1-273_ reveal distributions corresponding to the binding of one and two 4×-GAUGG oligonucleotides. Only one additional distribution is observed for N_1-220_ or N_1-273_ incubated with 4×-GAGAA oligonucleotide which corresponds to one oligonucleotide bound, (Figures S8-9, Table S3) confirming the preference for the 4×-GAUGG sequence for all N-terminal constructs.

**Figure 3.**
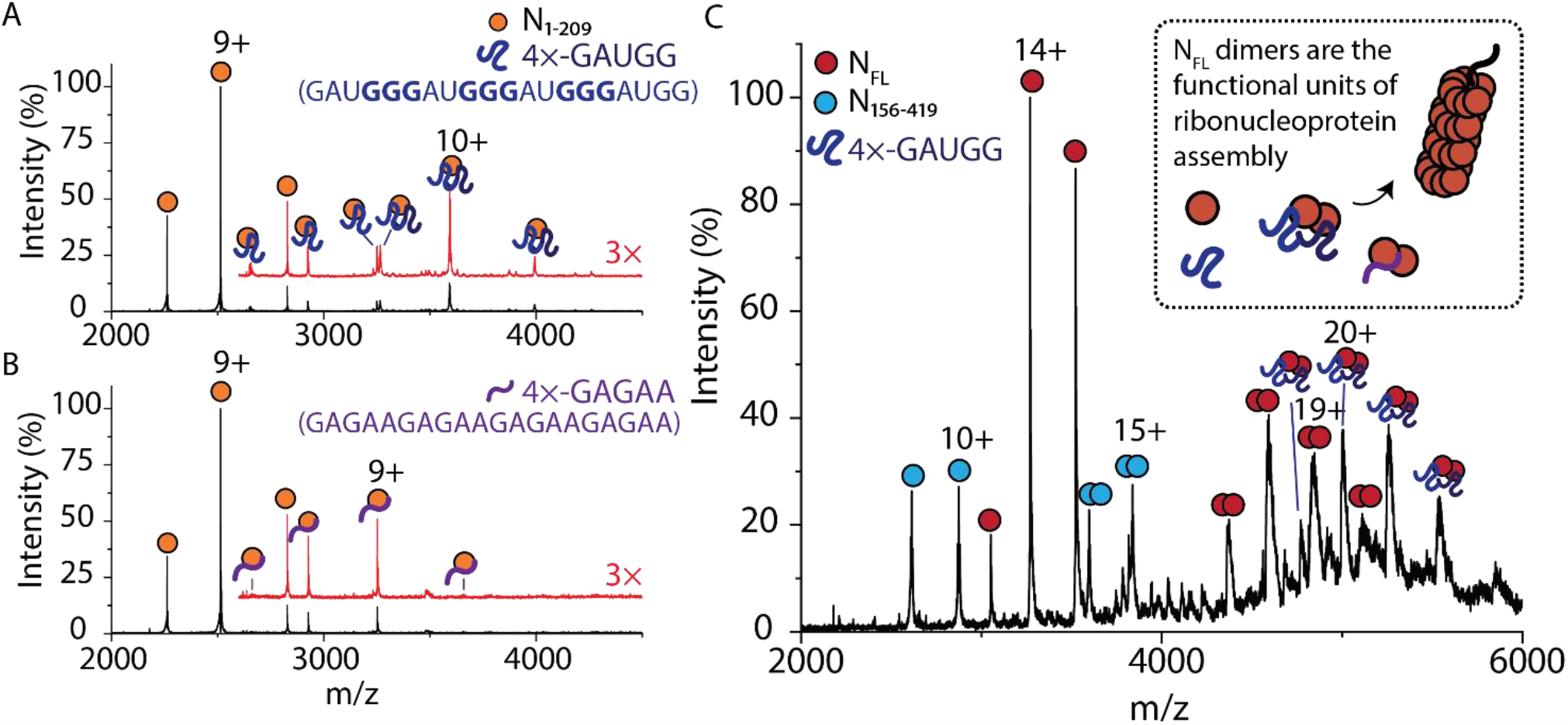
The RNA sequence influences binding stoichiometry to N protein. (A) Mass spectrum of N_1-209_ after incubation with 4×-GAUGG RNA oligonucleotides in a molar ratio of 1:4. Two additional charge state distributions are observed that correspond to one and two RNA oligonucleotides bound to N_1-209_. The mass spectrum at *m/z* >2700 was magnified 3× and offset for clarity (red trace). (B) Mass spectrum of N_1-209_ after incubation with 4×-GAGAA RNA oligonucleotides in a molar ratio of 1:4. One additional charge state distribution is observed that corresponds to one RNA oligonucleotide bound to N_1-209_. The mass spectrum at *m/z* >2700 was magnified 3× and offset for clarity (red trace). (C) Mass spectrum of N_FL_ after incubation with 4×-GAUGG RNA oligonucleotides in a molar ratio of 1:4. Monomers and dimers of N_FL_ (red circles) and N_156-419_ (blue circles) are observed. An additional peak series between 4300 and 5600 *m/z* corresponds to two 4×-GAUGG RNA oligonucleotides bound to N_FL_ dimer. The scheme in the inset of (C) depicts N_FL_ dimer bound to RNA as the functional unit of ribonucleoprotein assembly.

To examine if this preference was also observed for full-length N, we incubated the protein with 4×-GAUGG oligonucleotide. A series of peaks was identified corresponding to the N protein dimer bound to two 4×-GAUGG oligonucleotides (Figure 3C). Similarly, in the presence of the 4×-GAGAA RNA oligonucleotide, an additional RNA-bound distribution is observed, however the deconvoluted mass indicates that only one 4×-GAGAA oligonucleotide is bound to the N_FL_ dimer (Figure S10, Table S3). No RNA binding to the monomeric form of N_FL_ is observed, regardless of oligonucleotide sequence. Considering the different properties of the two RNA oligonucleotides, the 4×-GAUGG oligonucleotide contains three GGG motifs which form short stem-loops and are known to contribute additively to the efficiency of genome packaging in related viruses.^22^ Furthermore, selective RNA packaging has been described as a feature of innate immune response evasion.^23^ Our results emphasize that RNA sequence, and likely the secondary structure, is important for interactions with the N-protein. Notably, only N_FL_ dimer binds RNA, suggesting that the dimer is the functional unit of the SARS-CoV-2 ribonucleoprotein assembly. The preference for the RNA sequence known to form stem loops also has implications in efficient genome packaging and likely contributes to an optimized packing density in the intact virion.

### N proteoforms interact directly with cyclophilin A

Cyclophilin A (CypA), a highly abundant immunophilin found in host cells, has been implicated in the replication cycle of coronaviruses^24^ and plays multifunctional roles in modulating immune responses. We sought to determine if CypA plays a role in SARS-CoV-2 infection through monitoring direct interactions of CypA with N_FL_ or N proteoforms. We incubated N_1-209_ and CypA in a 1:1 molar ratio and used native mass spectrometry to measure possible interactions. The mass spectrum reveals charge state distributions that correspond to monomeric N_1-209_ and CypA, and three distinct charge state distributions at *m/z* >3000 (Figure 4A). The three higher-*m/z* distributions correspond to: (i) heterodimers of N_1-209_ and CypA, (ii) homodimers of CypA, and (iii) a low population of homodimers of N_1-209_ (Table 2). We observe similar interactions for N_1-273_ (Figure S11). Notably, we do not observe evidence of CypA interacting with N_1-220_ or N_FL_ under the same conditions (Figure S12-14).

**Table 2.** Deconvoluted and sequence masses for N proteoform complexes

| **Protein/Complex** | **Deconvoluted Mass<sup>a</sup> ± s.d. (Da)** | **Expected Mass (Da)** |
| --- | --- | --- |
| 4x-GAUGG RNA | -- | 6,622.00 |
| 4x-GAGAA RNA | -- | 6,650.20 |
| N<sub>1-209</sub> + one 4x-GAUGG RNA | 29,233 ± 0.4 | 29,234.71 |
| N<sub>1-209</sub> + two 4x-GAUGG RNA | 35,917 ± 10 | 35,856.71 |
| N<sub>1-209</sub> + one 4x-GAGAA RNA | 29,262 ± 1 | 29,262.91 |
| N<sub>FL</sub> dimer + two 4x-GAUGG RNA | 105,131 ± 60 | 104,783.66 |
| Cyclophilin A (CypA) | 20,084 ± 0.4<br>20,220 ± 0.8 | 20,175.82 |
| CypA dimers | 40,373 ± 62 | 40,351.64 |
| N<sub>1-209</sub> + CypA | 42,867 ± 15 | 42,696.71 |
| N<sub>1-209</sub> dimers | 45,356 ± 15 | 45,225.42 |
| Cyclosporin A | 1,202.85 ± 0 | 1,202.63 |
| CypA + CsA | 21,287 ± 0.5 | 21,286.63 |
| Antibody (monomer) | 137,990 ± 125<br>145,193 ± 109 | -- |
| N<sub>FL</sub> + antibody | 183,931 ± 102 | 183,759<br>190,962 |
| 2 N<sub>FL</sub> + antibody | 229,984 ± 36 | 229,528<br>236,731 |

<sup>a</sup> Determined using at least three adjacent charge states

**Figure 4.**
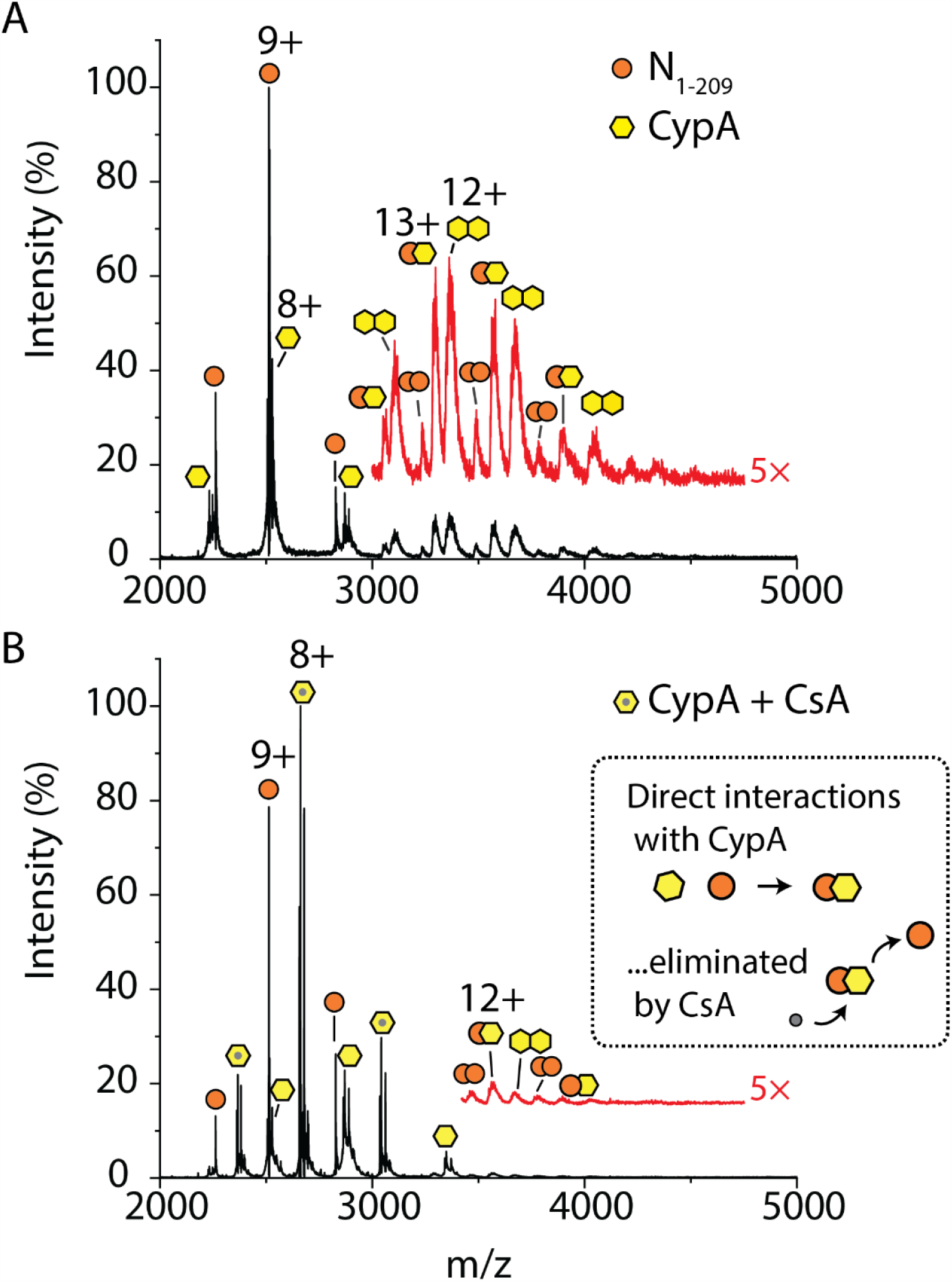
N proteoforms directly interact with cyclophilin A. (A) Mass spectrum of N_1-209_ after incubation with cyclophilin A (CypA) in a 1:1 molar ratio. The mass spectrum at *m/z* >3000 was magnified 5× and offset for clarity (red trace). Three charge state distributions that correspond to a low population of homodimers of N_1-209_, homodimers of CypA, and heterodimers of N_1-209_-CypA. (B) Mass spectrum of N_1-209_ after incubation with CypA and cyclosporin A (CsA) in a molar ratio of 1:1:2. CypA preferentially binds CsA as demonstrated by the charge state distribution centered at 8+ which corresponds to the CypA-CsA complex. The mass spectrum at *m/z* >3500 was magnified 5× and offset (red trace) to highlight the exceedingly low abundance of N_1-209_-CypA heterodimers that persist after CsA treatment. The scheme (inset of B) depicts CsA competitively binding to CypA and abolishing the N_1-209_-CypA interaction.

To determine if the interaction between N_1-209_ and CypA could be inhibited by an approved immunosuppressant cyclosporin A (CsA), we incubated the N_1-209_:CypA-complex with a 2-fold molar excess of the drug (Figure 4B). The mass spectrum reveals three abundant charge state distributions: (i) a distribution centered at 9+ corresponding to N_1-209_ monomers, (ii) a highly abundant distribution centered at a charge state of 8+ that corresponds to CypA bound to CsA, and (iii) a low abundant distribution that corresponds to monomeric CypA (Table 2). Very low abundance distributions of N_1-209_ homodimers, CypA homodimers, and N_1-209_-CypA heterodimers are barely detected following magnification >3000 *m/z*. Therefore, we can conclude that the 1:1 interaction between CypA and the N_1-209_ proteoform can be inhibited by CsA binding to CypA.

### Antigenic regions of N are located at the C-terminus

Since N protein is detected by antibodies with higher sensitivity than any other structural proteins of SARS-CoV-2^4^ we sought to determine if N proteoforms shared similar immunogenic properties as N_FL_. We first incubated N_FL_ with a monoclonal antibody (mAb) raised against the full-length protein in molar ratios of 1:1 (Figure 5A). The mass spectrum reveals three distributions >6000 *m/z* with deconvoluted masses that correspond to the apo antibody as well as one and two N_FL_ bound (Table 2, Figure S15). When the same mAb was incubated with three N-terminal proteoforms (N_1-209_, N_1-220_, and N_1-273_) we did not observe any mAb binding in the mass spectrum (Figure 5B). To validate these results with conventional methods we turned to protein detection by Coomassie stain and immunoblotting (Figure 5B inset). The Coomassie stain detects N_FL_ and all proteoforms in high abundance as indicated by the dark blue bands. In contrast, the immunoblot reveals dark bands for only N_FL_ and N_156-419_, including higher order oligomers of N_156-419_ that were not detected by Coomassie staining or mass spectrometry. N_1-209_ and N_1-273_ are barely detectable by the mAb, and N_1-220_ completely evades mAb detection. This suggests that N_1-209_, N_1-220_, and N_1-273_ may go undetected in viral infection. If this were the case, then we would expect to see a lowered antibody response to these N-terminal fragments in convalescent plasma from patients with COVID-19.

**Figure 5.**
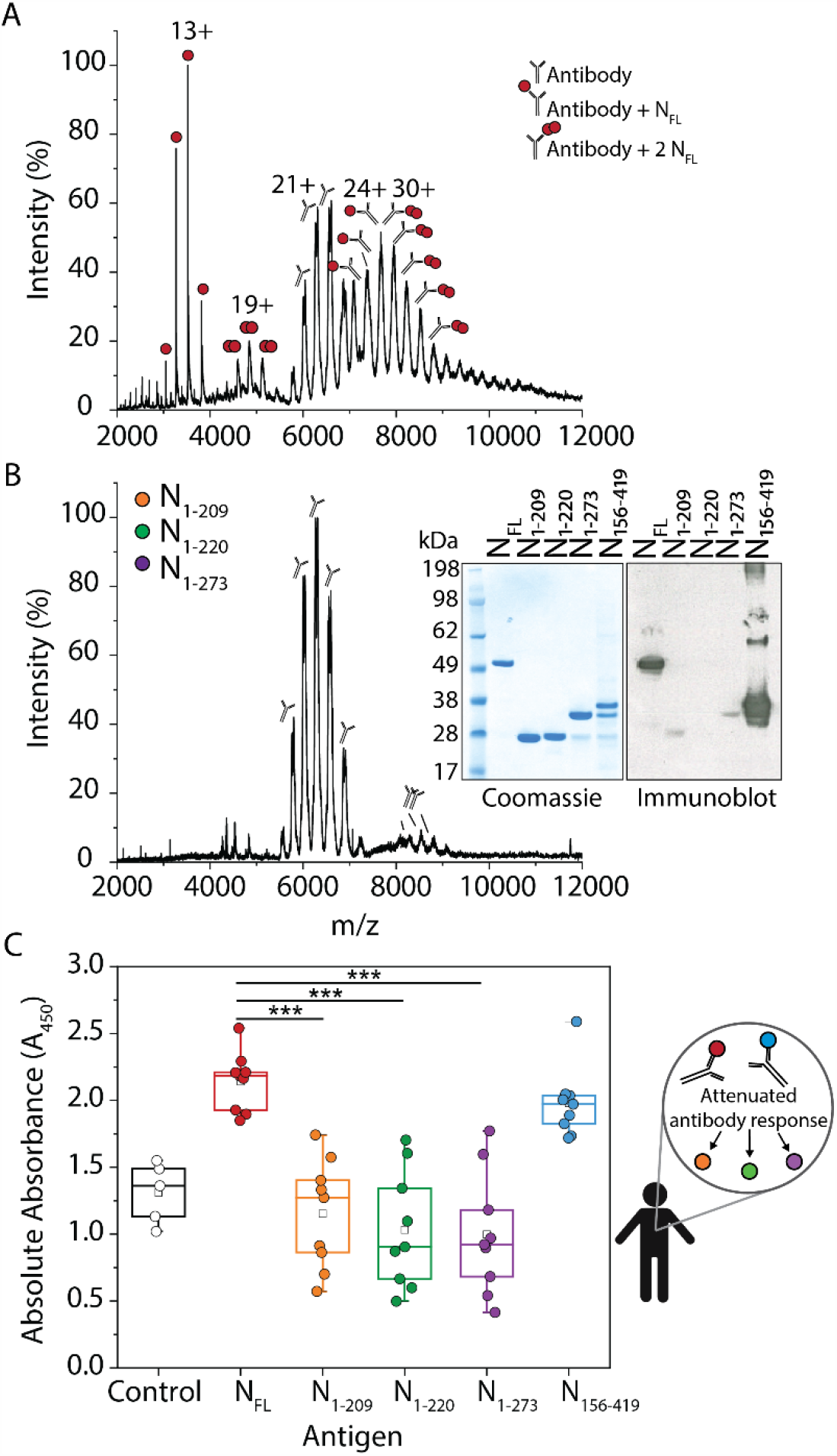
N_FL_ and N proteoforms have distinct immunological roles. (A) Mass spectrum of N_FL_ after incubation with a monoclonal antibody raised against the full-length N protein in a molar ratio of 1:1. We observe five charge state distributions that correspond to N_FL_ monomers (centered at 13+), N_FL_ dimers (centered at 19+), monomeric antibody (centered at 21+), antibody bound to one NFL (centered at 24+), and a population of antibody bound to N_FL_ dimer (centered at 30+). (B) Mass spectrum of a mixture of N_1-209_, N_1-220_, and N_1-273_ incubated with a monoclonal antibody in a molar ratio of 1:1:1:1. Charge state distributions are observed for antibody monomers and dimers. No binding to the antibody is observed for N proteoforms. This result was confirmed by immunoblot (see inset). SDS-PAGE shows that N_FL_, N_1-209_, N_1-220_, N_1-273_, and N_156-419_ are detected in high abundance by Coomassie stain. The same proteins analyzed by immunoblot show that only N_FL_ and N_156-419_ are detected by the antibody. (C) The box- and whisker plot depicts the antibody response to N_FL_ and N proteoforms using plasma from nine patients collected > 6 months following initial COVID-19 diagnosis. The antibody response was determined using the absolute absorbance following colorimetric detection of a sandwich ELISA where the immobilized antigen was N_FL_, N_1-209_, N_1-220_, N_1-273_, or N_156-419_. The control represents the measured response for a monoclonal antibody raised against the full-length N protein bound to N_FL_. The squares represent the mean, the center line represents the median, and the box represents the first quartile (25-75%) of the distributed data. Asterisks represent statistically significant differences when compared to N_FL_; p-values for N_1-209_, N_1-220_, and N_1-273_ are 8.25e^-6^, 4.18e^-6^, and 4.27e^-6^, respectively. The antibody response for N_156-419_ compared to NFL was not statistically different with a p-value of 0.17.

To test this hypothesis we obtained convalescent plasma from nine patients > 6 months after an initial diagnosis of COVID-19 and studied the antibody response to the N proteoforms characterized herein. The experiment was carried out using an enzyme-linked immunosorbent assay (ELISA) using all five proteoforms (N_FL_, N_1-209_, N_1-220_, N_1-273_, N_156-419_) as the antigen to which antibodies were captured. The plasma antibodies were “sandwiched” using an anti-IgG detection antibody conjugated with horseradish peroxidase for colorimetric detection. The antibody response for all nine patients was measured as a function of the absorbance and displayed as a box plot (Figure 5C). Using the mAb bound to N_FL_ as a positive control, we qualitatively concluded that all proteoforms react with antibodies present in convalescent plasma. However, when compared to N_FL_, the N-terminal proteoforms (N_1-209_, N_1-220_, N_1-273_) resulted in a significantly attenuated antibody response. Interestingly, the antibody response for N_FL_ and N_156-419_ were not statistically different, suggesting that the antigenic site for antibody recognition is localized to the C-terminus of the N protein. That this preferential antibody response is still detectable six months after COVID-19 diagnosis implies that N_156-419_ is an attractive target for vaccine design against SARS-CoV-2.

## Conclusion

We present a comprehensive characterization of SARS-CoV-2 N protein and highlight potential features that might influence SARS-CoV-2 infectivity (Figure 6). Specifically, we find that N protein undergoes proteolysis at highly conserved residues in the vicinity of the linker region, separating the two major domains (the RNA binding and oligomerization domains). We identify various stoichiometries of N proteoforms that are influenced by pH, explicitly N_156-419_, which forms stable oligomers under both high and low pH solution conditions. We also show that N_FL_ and N proteoforms bind RNA with a preference for GGG-motifs and present evidence for N_FL_ dimers being the functional unit of assembly in ribonucleoprotein complexes. Furthermore, we determined that immunophilin CypA binds directly to N_1-209_ and N_1-273_, but not N_FL_ or N_1-220_, an interaction that can be inhibited through addition of the immunosuppressant cyclosporin A. To test the immunogenicity of N proteoforms, we used a recombinant antibody and immunoblot techniques to demonstrate that the antigenic site of SARS-CoV-2 N protein resides towards the C-terminus. To test this in an *in vivo* scenario we obtained convalescent plasma from nine patients > 6 months after initial COVID-19 diagnosis. We discovered a heightened response for the N_FL_ and N_156-419_ relative to N-terminal proteoforms N_1-209_, N_1-220_, and N_1-273._

**Figure 6.**
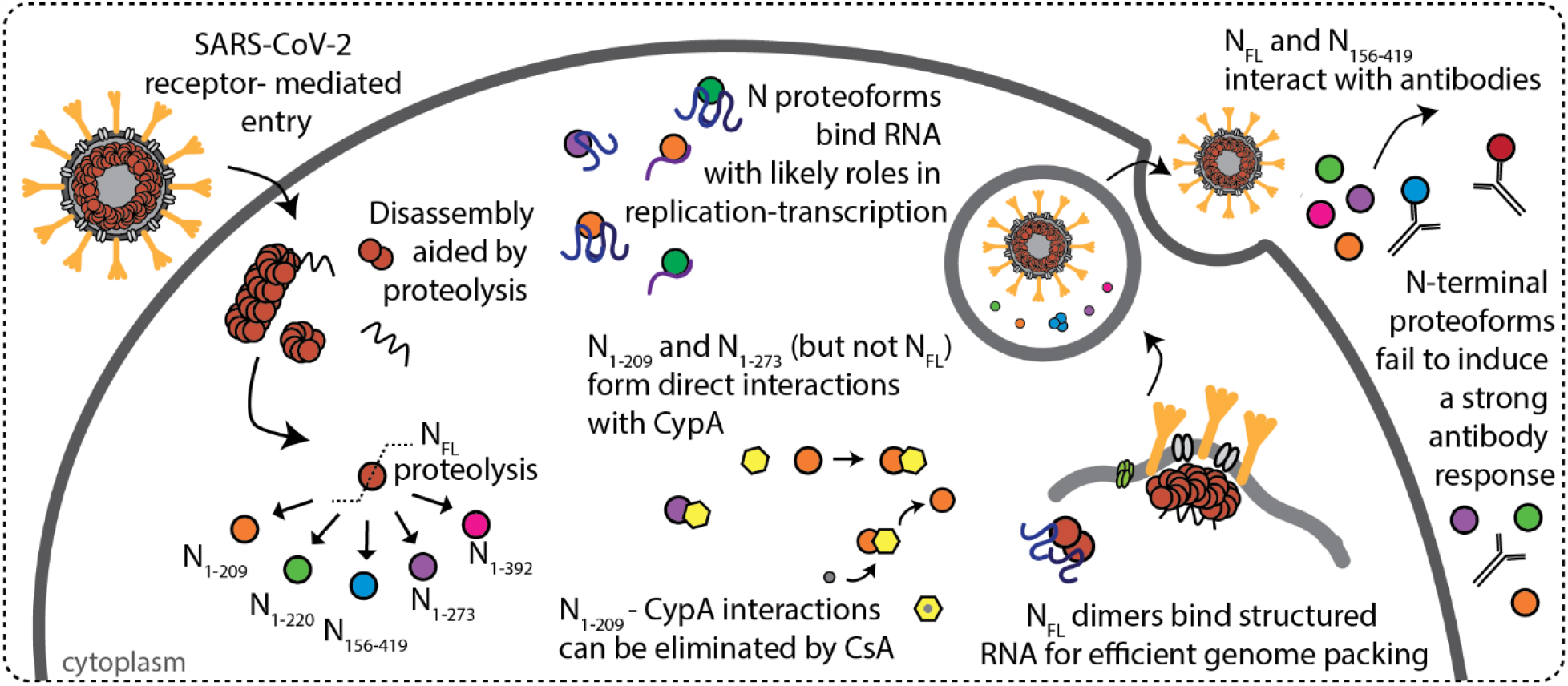
Scheme depicting features of SARS-CoV-2 N protein during infection. N protein undergoes proteolysis in a highly specific fashion to produce N_1-209_, N_1-220_, N_1-273_, N_156-419_, and N_1-392_. N_FL_ and N proteoforms bind RNA with a preference for structured RNA and N_FL_ dimers are likely functional unit of assembly in ribonucleoprotein complexes. Immunophilin CypA binds directly to N_1-209_ and N_1-273_, but not N_FL_ or N_1-220_, and the interaction can be inhibited through addition of cyclosporin A (CsA). N_156-419_ and N_FL_ interact with antibodies from convalescent plasma, while proteoforms N_1-209_, N_1-220_, and N_1-273_ fail to induce the same antibody response.

In general, proteolysis leads to changes in protein function and is a key strategy in viral proliferation.^25^ The nucleocapsid protein of SARS-CoV, responsible for the SARS pandemic of 2002, was shown to undergo similar proteolytic cleavage, however the precise role of proteolysis in SARS-CoV N protein remains elusive.^26,27,28^ New evidence reveals proteolytic cleavage of multiple viral proteins during SARS-CoV-2 infection, including extensive cleavage of the N protein.^29^ Our results share striking similarities with the cleavage sites observed for both SARS-CoV and N protein from SARS-CoV-2 infected cells^29^ and therefore elucidating the role of N proteoforms represents a necessary endeavor for targeting cellular processes involved in viral proliferation.

One of the most critical steps of viral proliferation is the packaging of genomic material and its assembly into new virions. Genome packaging of RNA viruses is highly selective and depends on specific nucleotide sequences and complex structural elements called packaging signals (Psi).^30^ It has recently been demonstrated that HIV polyproteins have a stronger tendency to oligomerize in complex with Psi-RNA relative to non-Psi RNA.^31^ Our results indicate that N_FL_ and N proteoforms exhibit preference for RNA sequences that mimic key structural features of the genomic RNA.^32^ We anticipate that the more efficient binding of N protein dimers to RNA with GGG-motifs underlies the selective packaging of genomic RNA in SARS-CoV-2. The disruption of structural features of viral RNA has been proven to inhibit replication in some viruses.^33^ In the case of SARS-CoV-2, disruption of preferred RNA structures, and therefore disruption of N protein-RNA interactions, presents a promising strategy to intervene in replication and the potential to develop live attenuated vaccines.

Immunological roles of N proteoforms have been demonstrated by the highly specific interactions between cyclophilin A and the N_1-209_ and N_1-273_, but not N_FL_. Cyclophilin A has been found in mature virions of HIV and is known to play a key role in HIV replication. Interestingly, the interaction between cyclophilin A and the HIV capsid protein is known to be conformation-dependent^34^; we anticipate such parallels with SARS-CoV-2 and we can only speculate that N_1-220_ does not interact with CypA because of a conformational change not exhibited by the other proteoforms. In addition to its role in virus replication, cyclophilin A is implicated in inflammation and is a mediator of cytokine production.^35^ In cases of severe COVID-19 infection, patients have presented with aberrant inflammatory responses, known as “cytokine storms” which can be fatal.^36^ The symptoms identifying cytokine storms have been established,^37^ however, the viral component(s) that trigger such a response have yet to be elucidated. We can only speculate that specific interactions between N proteoforms and cyclophilin A, as demonstrated here, may play a role in producing such an aberrant immune response. Pathways to intervene with such interactions could therefore prove beneficial as a component of treatment strategies.

Considering antibody responses, proteoforms N_1-209_, N_1-220_, and N_1-273_ fail to activate the same antibody response as the full-length protein. These proteoforms could thereby contribute to virtually unchecked virus proliferation, depending on their functional roles. By contrast, N_156-419_ produced an antibody response from convalescent plasma that could not be differentiated from the full-length protein. This result strongly implies that that the antigenic site of the N protein is localized to the C-terminus which is in accord with epitope mapping of the homologous SARS-CoV N protein.^38,39,40,41^ We speculate that the oligomerization of antigenic N_156-419_, which readily forms dimers, trimers and hexamers as a function of pH, may serve to increase the avidity to antibodies. We propose that N_156-419_ oligomers may act as decoys for neutralizing antibodies. This proposal is in line with reports for a number of viruses, where antigenic proteins, or protein complexes, are shed in high abundance to consume neutralizing antibodies.^42,43,44^ This strategy has also been used in the development of antiviral therapies for SARS-CoV-2 wherein soluble variants of the host receptor (decoy receptors) are used to bind and neutralize the infectious virus.^45^

Several additional therapeutic avenues are highlighted by this study. Firstly, inhibition of the proteolysis reaction would prevent the formation viral proteoforms that may be fine-tuned for various functions within the viral lifecycle. While the mechanism behind the proteolysis of N-protein is not yet understood, systematic mutation of cleavage sites defined here could lead to mechanistic and structural insights to enable small molecule screening of potential protease inhibitors. Knowledge of N protein-structured RNA interactions could also aid the design of new therapeutics that would inhibit successful replication. However, since N protein oligomerizes in the absence of RNA,^46^ *de novo* drug design of assembly inhibitors is complex as it is necessary to consider oligomerization propensity of the various proteoforms at the pH regimes encountered in the cellular environment. CsA and cyclosporine-derivatives, such as Alisporivir, however are more straightforward to track and have become attractive candidates to treat COVID-19.^47^ Disruption of the cyclophilin A-N-proteoform interactions shown here provides a convenient means of screening potential inhibitors for hit to lead optimization.

Finally, although we have uncovered many potential roles of N protein and its proteoforms, interactions with at least nine host proteins involved in RNA processing have yet to be defined.^12^ Furthermore, defining post-translational modifications,^48^ and interactions with other host proteins is a necessary endeavor that will provide a more complete knowledge of the multifaceted roles of SARS-CoV-2 N protein. The results presented here however do prompt further investigation of a number of therapeutic strategies including inhibition of the proteolyis reaction, perturbation of N-protein RNA binding, prevention of cyclophilin A:proteoform interactions, and possible development of N_156-419_ as a vaccine candidate. Given the likelihood that no one intervention is likely to ameliorate the complex symptoms of COVID-19 infection, our findings contribute proteoform-specific information that may guide some of the many therapies currently under investigation.

## Materials and Methods

### Ethics

Patients were recruited from the John Radcliffe Hospital in Oxford, United Kingdom, between March and May 2020 with written and informed consent. Participants were identified from hospitalization during the SARS-COV-2 pandemic and recruited into the Sepsis Immunomics and International Severe Acute Respiratory and Emerging Infection Consortium World Health Organization Clinical Characterisation Protocol UK (IRAS260007 and IRAS126600). Patient samples were collected at least 28 days from the start of their symptoms. Ethical approval was given by the South Central–Oxford C Research Ethics Committee in England (reference: 13/SC/0149), Scotland A Research Ethics Committee (reference: 20/SS/0028) and World Health Organization Ethics Review Committee (RPC571 and RPC572l; 25 April 2013).

### Plasmid construction and cell growth

A codon-optimized synthetic gene corresponding to the full length nucleocapsid protein (Thermo GeneArt, Regensburg, Germany) was cloned into a modified pET28a vector using the In-Fusion cloning kit (Takara Bio Saint-Germain-en-Laye, France). The resulting plasmid encoded for an N-terminal His_6_ tag followed by thrombin and tobacco etch virus cleavage sequences upstream of the full-length nucleocapsid protein sequence. To generate the nucleocapsid proteoforms, the desired sequences pertaining to the truncated forms of the N protein were subcloned from the synthetic gene using polymerase chain reaction (Phusion polymerase, New England Biolabs, Hertfordshire, UK). All genes were cloned into the modified pet28 vector and gene sequences were confirmed by Sanger Sequencing.

Plasmids were transformed into BL21 (DE3) and streaked onto LB agar plates supplemented with 50 mg mL^-1^ kanamycin. Several colonies were used to inoculate 100 mL of LB broth supplemented with kanamycin and grown at 37 °C overnight. 10 mL of the overnight precultures were used to inoculate 1 L of LB broth supplemented with kanamycin. Cell cultures were grown to OD_600_ ~ 0.6 before inducing protein expression with 0.5 µg mL^-1^ IPTG. Cells were grown for an additional 4 hours at 37 °C before harvesting via centrifugation (5000 x g, 10 minutes, 4 °C). Cell pellets were flash frozen in liquid nitrogen and stored at −80 °C until use.

### Protein Purification

Cell pellets were resuspended in lysis buffer (25 mM Tris-HCl pH 8.0, 500 mM NaCl, 5 mM MgCl_2_, 5 mM β-mercaptoethanol, 5 mM imidazole, 10% v/v glycerol) containing EDTA-free protease inhibitor tablets (Roche). Cells were lysed by five passes through a microfluidizer (prechilled to 4 °C) at 20,000 psi. Cell debris was pelleted by centrifugation (20,000 x g, 20 minutes, 4 °C). The supernatant containing soluble nucleocapsid protein was passed through a 0.45 µm filter.

Supernatant was loaded onto a Ni-NTA column pre-equilibrated in loading buffer (25 mM Tris-HCl pH 8.0, 500 mM NaCl, 5 mM MgCl2, 5 mM β-mercaptoethanol, 20 mM imidazole, 10% v/v glycerol) and allowed to pass via gravity flow. To remove common contaminating proteins, a heat-treated BL21 (DE3) *E. coli* lysate in loading buffer containing 10 mM MgATP was passed over the immobilized nucleocapsid protein using a protocol by Rial and Ceccarelli.^49^ The resin was washed with 10 column volumes of wash buffer (25 mM Tris-HCl pH 8.0, 500 mM NaCl, 5 mM MgCl2, 5 mM β-mercaptoethanol, 80 mM imidazole, 10% v/v glycerol), then eluted twice with 10 mL of elution buffer (25 mM Tris-HCl pH 8.0, 500 mM NaCl, 5 mM MgCl_2_, 5 mM β-mercaptoethanol, 400 mM imidazole, 10% v/v glycerol).

The eluted protein was mixed with TEV protease in a 100:1 (w/w) and loaded into a 3 kDa MWCO dialysis cassette (Thermo Fisher Scientific, United Kingdom). Cleavage of the His_6_-thrombin-TEV tag was carried out overnight at 4 °C in lysis buffer. The cleaved tag and TEV protease were separated from the untagged protein using reverse immobilized metal affinity chromatography on a Ni-NTA column prepared in loading buffer. The flow-through containing the untagged protein was collected and concentrated in a 10k MWCO centrifugal filter before MS analysis. Protein concentration was determined using UV-VIS spectroscopy by monitoring the absorbance at 280 nm with a theoretical extinction coefficient (ε ~ 43890 M^-1^ cm^-1^) determined using the Expasy Protparam tool.

### Size Exclusion Chromatography

Full-length N protein and N proteoforms were separated using a Superdex 10/300 increase GL column equilibrated in lysis buffer. Fractions corresponding to N_FL_ and N proteoforms were pooled and concentrated to ~10 µM before MS analysis.

### Native Mass Spectrometry

RNA oligonucleotides of repeating sequences (4×-GAUGG, 4×-GAGAA) were purchased from Integrated DNA Technologies. Recombinant human cyclophilin A (product ab86219) was purchased from Abcam (Cambridge, United Kingdom). Cyclosporine A purchased from Merck Life Science (Dorset, United Kingdom). N protein and all binding partners were buffer exchanged or diluted into 500 mM NH_4_OAc pH 5.0, 7.4, or 8.0. Buffer exchange was carried out using Zeba™ Spin Desalting Columns, 7K MWCO (Thermo Fisher Scientific, United Kingdom).

Measurements were carried out on Qexactive UHMR or Orbitrap Eclipse. The Q-exactive instrument was operated in the positive ion mode using the manufacturer’s recommended parameters for native MS. The instrument was operated at a resolving power of 12,500 (at *m/z* 200). An electrospray was generated by applying a slight (~0.5 mbar) backing pressure to an in-house prepared gold-coated electrospray capillary held at ~1.2 kV relative to the instrument orifice (heated to ~100 °C).

An Eclipse Tribrid instrument was also used for native MS and top-down sequencing. The instrument was set to intact protein mode at standard ion routing multipole pressure of 10 mTorr. Ion voltages were set to transmit and detect positive ions at a resolving power of 12,500 (at *m/z* 200). An electrospray voltage of ~1.2 kV and ~0.5 mbar backing pressure were used for ion formation; desolvation was assisted using an instrument capillary temperature of ~100 °C. To identify the accurate mass and sequence of each proteoform, we used a similar approach to that described by Huguet et al.^17^ Briefly, a desired signal was isolated using the ion trap (10 *m*/*z* isolation window, charge state set to 10) and subjected to: (i) electron transfer dissociation (ETD, 3 ms activation time, 1.0×10^6^ ETD reagent target) to generate a charge-reduced series for accurate intact mass determination, and (ii) higher energy collisional dissociation (HCD) using ~30 – 50 V HCD collision energy to generate fragment ions.

Monoisotopic masses of fragment ions generated by HCD having a normalized intensity of 10% or higher were fed into Prosite Lite software.^50^ A series of candidate sequences that best matched the measured intact masses for each proteoform were generated. The monoisotopic fragment masses were matched to expected ions generated in silico based on the provided candidate sequence. Comparison of the statistical likelihood for each match compared to a series of candidate sequences (Table S2) localized the cleavage sites to those outlined in Figure 2D.

### Liquid Chromatography and Bottom-up Mass Spectrometry

Full-length N was separated from the lower molecular weight proteoforms on a 0.8 mm 4-12% bis-tris SDS-PAGE gel (Invitrogen) and stained with Coomassie Blue. Bands were excised from the gel, minced, and digested with sequencing grade trypsin (Promega, Madison, WI, U.S.A) at 37 °C overnight, extracted with 80% acetonitrile (0.1% formic acid), and dried on a vacuum concentrator. The extracted peptides were resolubilized in buffer A (H_2_O, 0.1% FA) and loaded onto a reverse phase trap column (Acclaim PepMap 100, 75µm x 2 cm, nano viper, C18, 3 µm, 100 Å, ThermoFisher, Waltham, MA, U.S.A.) using an Ultimate 3000 for 50 µL at a flow rate of 10 µL min^-1^. The trapped peptides were then separated using a 15 cm reverse phase analytical column (350 µm x 75 µm) packed in-house (3 µm C18 particles) using a 60 min linear gradient from 5% to 40% buffer B (80% acetonitrile, 20% water, 0.1% formic acid) at a flow rate of 300 nL min^-1^. The separated peptides were then electrosprayed in the positive ion mode into an Orbitrap Eclipse Tribrid mass spectrometer (ThermoFisher, San Jose, CA, USA) operated in data-dependent acquisition mode (3 s cycle time). Precursor and product mass analysis occurred in the Orbitrap analyzer (120,000 and 60,000 resolving power at *m/z* 200, respectively). High intensity (threshold: 1.0 × 10^4^) precursors with charge state between z = 2 and z = 7 were isolated with the quadrupole (0.5 *m/z* offset, 0.7 *m/z* isolation window) and fragmented using higher energy collision induced dissociation (HCD collision energy = 30%). Additional MS/MS scans for precursors within 10 ppm were dynamically excluded for 30 s following the initial selection. MS/MS scans were collected using an automated gain control setting of 1.0 × 10^4^ or a maximum fill time of 100 ms. LC-MS data were searched against both the *E. coli* proteome manually annotated with the SARS-CoV-2 nucleocapsid protein sequence using MaxQuant v1.6.17.0.

### Western Blot

Recombinant antibody generated from the full-length SARS-CoV-2 nucleocapsid protein (product ab272852) and anti-human secondary antibody were purchased from Abcam. Proteins were resolved on a 4-12% Bis-tris gel using SDS-PAGE and transferred to a PVDF membrane (pore size 0.45 µm). The membrane was blocked in 5% milk in TPBS for 1 hour at RT, incubated with primary antibody in 1:2000 dilution into blocking buffer (also 1 hour, RT) washed with TPBS and incubated with secondary antibody (1:10000) in blocking buffer (also 1 hour, RT). The PVDF membrane was incubated with Horseradish peroxidase chemiluminescent substrate (Pierce ECL Western Blotting Substrate, Thermo Scientific) before detection on photographic film and developed by an X-ray film processor (Xograph Compact X4).

### Enzyme-Linked Immunosorbent Assay

Recombinant antibody generated from the full-length SARS-CoV-2 nucleocapsid protein (product ab272852) and goat anti-human IgG (ab97225) conjugated with horseradish peroxidase (HRP) purchased from Abcam. Nickel coated clear 96-well plates were purchased from Thermo Scientific (Thermo Fisher Scientific, United Kingdom). Plates came pre-blocked and 50 µg of his-tagged N proteoforms were loaded into each well and allowed to incubate at room temperature for 1 hour. Plates were washed three times with 200 µL of phosphate buffered saline (PBS) containing 0.05% Tween-20 (TPBS). Patient plasma was diluted 8-fold with PBS and 100 µL was added to each well and the controls were carried out using 20 ng of a monoclonal antibody raised against the full-length protein (ab272852) added to wells containing N_FL_. Antibodies from patient plasma and the control antibody were allowed to incubate overnight at 4 °C. Plates were washed three times with TPBS. Goat anti-human IgG secondary antibody was diluted 1:50,000 in PBS, 100 µL added to each well, and the plates were incubated at room temperature for 1 hour. Plates were washed a further 3 times with TPBS. Colorimetric detection was carried out using a TMB chromogenic substrate kit for HRP detection (Thermo Fisher Scientific, United Kingdom). The reaction was quenched after 5 minutes using 2 M sulfuric acid, resulting in a yellow color. Absorbance measurements were immediately carried out at 450 nm using a microplate reader (BMG Labtech, Aylesbury, United Kingdom).

## Safety Statement

No unexpected or significant safety hazards are associated with the reported work.

## Supporting information

Supporting Information for Proteoforms of the SARS-CoV-2 nucleocapsid protein are primed to proliferate the virus and attenuate the antibody response

## Acknowledgements

We thank Edward Emmott for helpful discussions and Alexander Mentzer, Gavin Screaton, Tao Dong, and Yanchun Peng for providing convalescent plasma from anonymized COVID-19 patients. We are grateful to the patients for donating their samples, and to the research teams involved in consent, recruitment and sampling of these participants. We are grateful for generous support provided by the University of Oxford COVID-19 Research Response fund and its donors (BRD00230). C.V.R. is also a part of the COVID-19 mass spectrometry consortium.^51^ Work in the C.V.R. laboratory is supported by a Medical Research Council (MRC) program grant (MR/N020413/1), a European Research Council Advanced Grant ENABLE (695511), and a Wellcome Trust Investigator Award (104633/Z/14/Z). C.A.L. is supported by the European Union’s Horizon 2020 research and innovation programme under the Marie Skłodowska-Curie grant agreement GPCR-MS 836073. T.J.E. is supported by the Royal Society as a Royal Society Newton International Fellow.

## Author Contributions

C.A.L, and T.J.E. designed the protein constructs; C.A.L., T.J.E., and J.R.B. expressed and purified protein constructs. C.A.L., T.J.E., and C.V.R., conceived the experiments; C.A.L. performed the experiments; T.J.E, J.R.B., and C.V.R supported the experiments; all authors co-wrote the paper.

## Ethics declarations

The authors declare no competing interests. Carol Robinson provides consultancy services for OMass Therapeutics.

## Figures and Legends

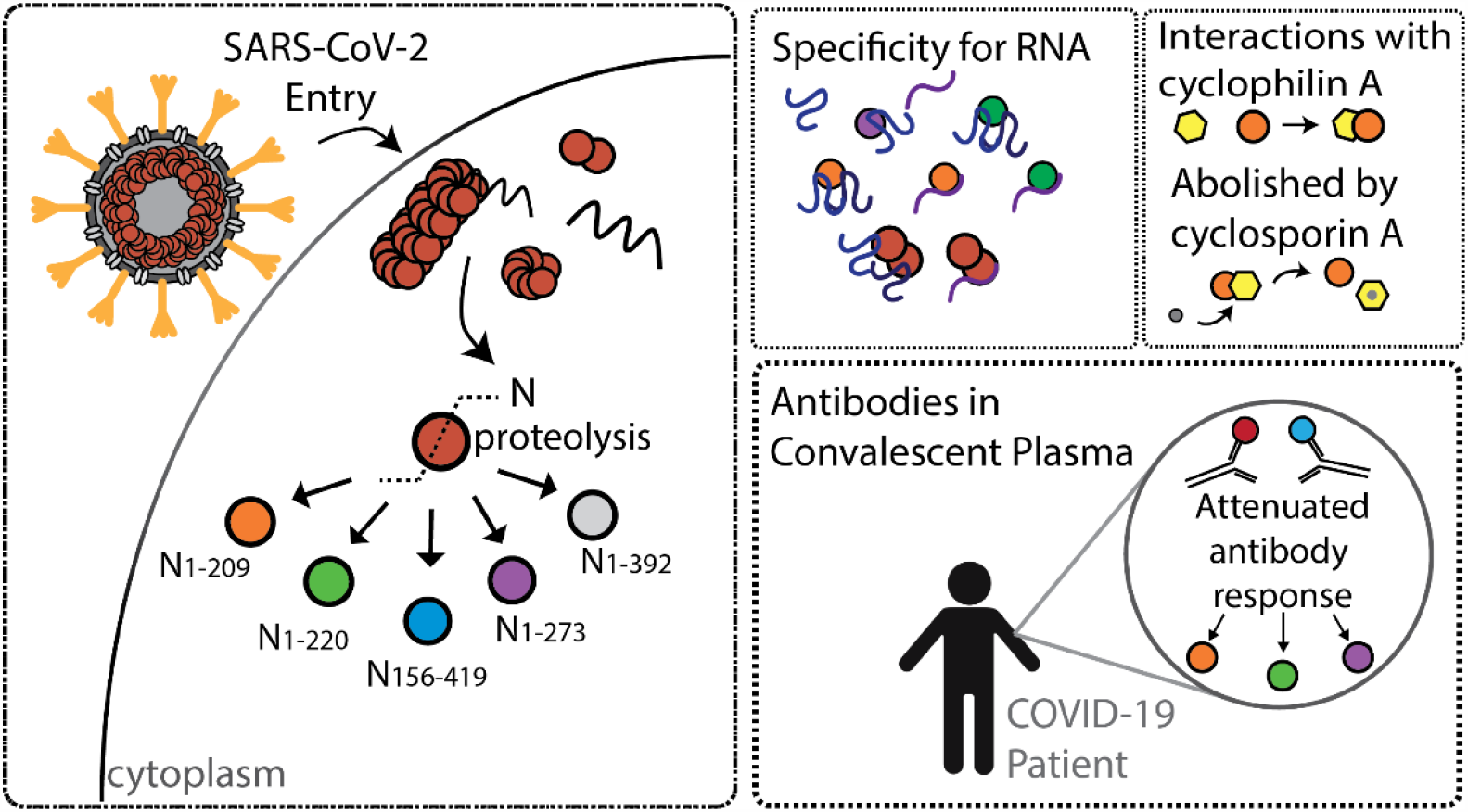

