## Supporting Information for Proteoforms of the SARS-CoV-2 nucleocapsid protein are primed to proliferate the virus and attenuate the antibody response for "Proteoforms of the SARS-CoV-2 nucleocapsid protein are primed to proliferate the virus and attenuate the antibody response"

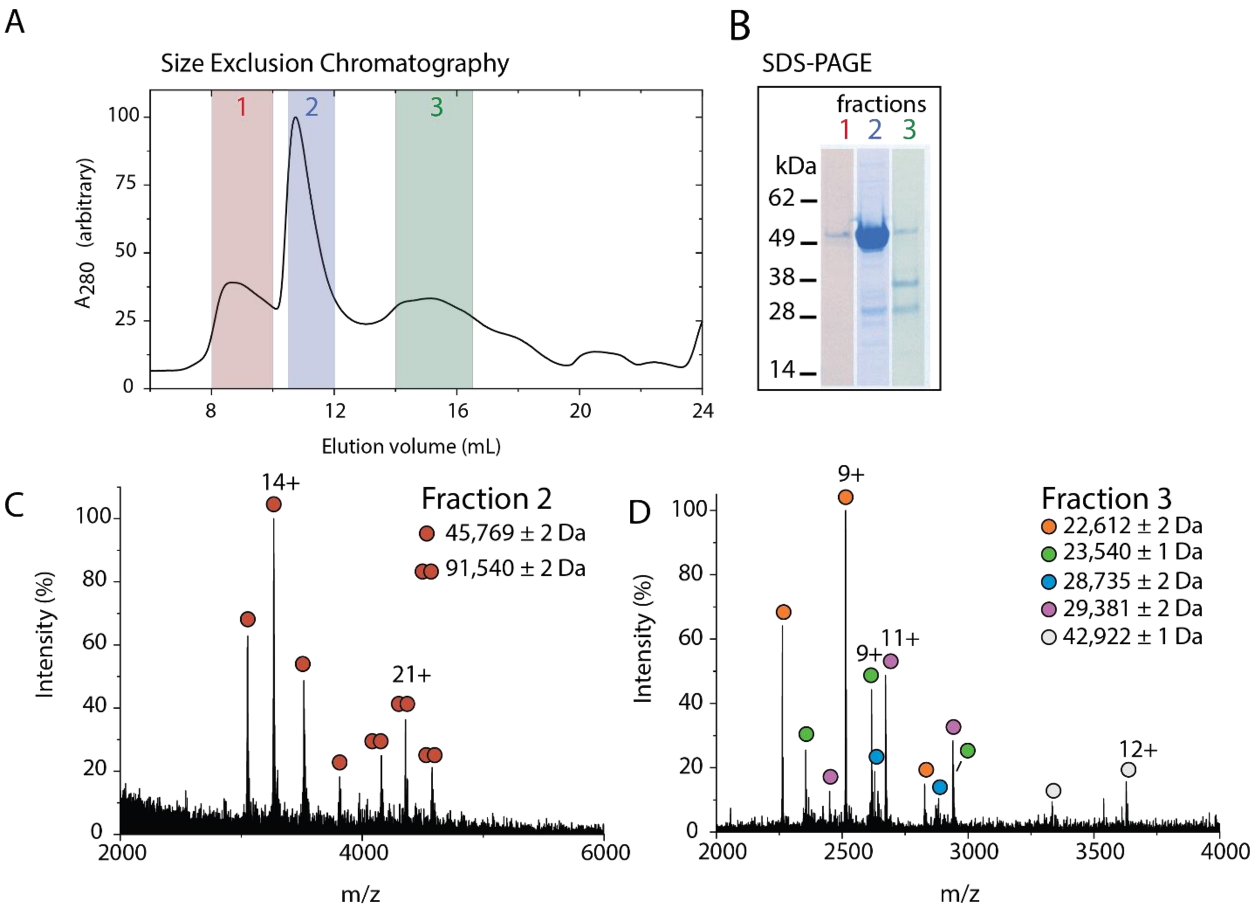


**Figure S1**. Separation of N proteoforms from $N_{FL}$ using size exclusion chromatography. (A) Purified N protein was separated using size exclusion chromatography and the absorbance (A280) was monitored throughout fractionation. The fractions from three distinct peaks (denoted 1, 2 and 3) were pooled and concentrated. (B) SDS-PAGE analysis of the pooled and concentrated fractions indicated that $N_{FL}$ was the dominant species in fraction 1 and 2, while fraction 3 contained mostly lower molecular weight proteoforms. (C) The mass spectrum of fraction 2 revealed monomers and dimers of full-length N. (D) The mass spectrum of fraction 3 revealed five proteoforms that range in mass from 22,612 to 42,922 Da. No interpretable mass spectrum was collected for fraction 1; the protein in fraction 1 eluted in the void volume of the column, indicating high molecular weight oligomers or aggregates.

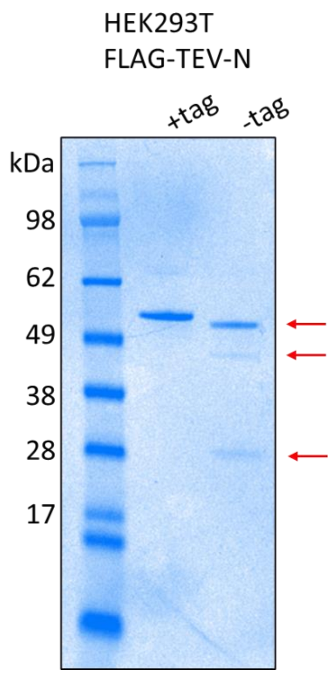


**Figure S2.** N protein expressed in mammalian cells undergoes proteolysis. N protein was expressed in human embryonic kidney (HEK293T) cells and the purified protein (pre- and post-tag removal) were analyzed by SDS-PAGE. Coomassie stain reveals three distinct bands that emerge after affinity tag removal of the pure protein.

**Table S1**. Statistical analysis for HCD fragmentation products matched to candidate sequences.

**Deconvoluted mass of charge state distribution**: 22,611 ± 1
**Tentative Assignment**: $N_{1\text{-}209}$

| Candidate Sequence | Sequence Mass (Da) | Precursor Charge State | Fragments matched | P-score | Coverage (%) |
|---|---|---|---|---|---|
| $N_{FL}$ | 45,769.83 | 8+ | 0 | 1 | 0 |
| | | 9+ | 10 | 1.40e-9 | 2 |
| | | 10+ | 0 | 1 | 0 |
| $N_{1\text{-}208}$ | 22,456.52 | 8+ | 0 | 1 | 0 |
| | | 9+ | 10 | 1.40e-9 | 5 |
| | | 10+ | 0 | 1 | 0 |
| **$N_{1\text{-}209}$** | **22,612.71** | 8+ | 0 | 1 | 0 |
| | | **9+** | **11** | **7.60e-11** | **5** |
| | | 10+ | 1 | 0.07 | 0 |
| $N_{1\text{-}210}$ | 22,743.9 | 8+ | 0 | 1 | 0 |
| | | 9+ | 9 | 2.20e-8 | 4 |
| | | 10+ | 0 | 1 | 0 |
| $N_{215\text{-}419}$ | 22,687.6 | 8+ | 0 | 1 | 0 |
| | | 9+ | 1 | 4.60e-1 | 0 |
| | | 10+ | 0 | 1 | 0 |
| $N_{216\text{-}419}$ | 22,572.51 | 8+ | 0 | 1 | 0 |
| | | 9+ | 1 | 4.60e-1 | 0 |
| | | 10+ | 0 | 1 | 0 |

N G S M S D N G P Q N Q R N A P R I T F G G P S D S 25
26 T G S N Q N G E R S G A R S K Q R R P Q G L P N N 50
51 T A S W F T A L T Q H G K E D L K F P R G Q G V P 75
76 I N T N S S P D D Q I G Y Y R R A T R R I R G G D 100
101 G K M K D L S P R W Y F Y Y L G T G P E A G L P Y 125
126 G A N K D G I I W V A T E G A L N T P K D H I G T 150
151 R N P A N N A A I V L Q L P Q G T T L P K G F Y A 175
176 E G S R G G S Q A S S R S S S R S R N S S R N S T 200
201 P G S S R G T S P A R C

**Deconvoluted mass of charge state distribution**: 23,540 ± 0.3
**Tentative Assignment**: $N_{1\text{-}220}$

| Candidate Sequence | Sequence Mass (Da) | Precursor Charge State | Fragments matched | P-score | Sequence Coverage (%) |
|---|---|---|---|---|---|
| $N_{FL}$ | 45,769.83 | 9+ | 5 | 3.00e-3 | 1 |
| | | 10+ | 0 | 1 | 0 |
| $N_{1\text{-}219}$ | 23,470.65 | 9+ | 5 | 3.00e-3 | 2 |
| | | 10+ | 0 | 1 | 0 |
| **$N_{1\text{-}220}$** | 23,541.73 | **9+** | **9** | **7.60e-7** | **4** |
| | | 10+ | 1 | 0.07 | 0 |
| $N_{1\text{-}221}$ | 23,654.89 | 9+ | 5 | 3.00e-3 | 2 |
| | | 10+ | 0 | 1 | 0 |
| $N_{206\text{-}419}$ | 23,499.52 | 9+ | 1 | 0.61 | 0 |
| | | 10+ | 0 | 1 | 0 |
| $N_{205\text{-}419}$ | 23,586.59 | 9+ | 1 | 0.61 | 0 |
| | | 10+ | 0 | 1 | 0 |

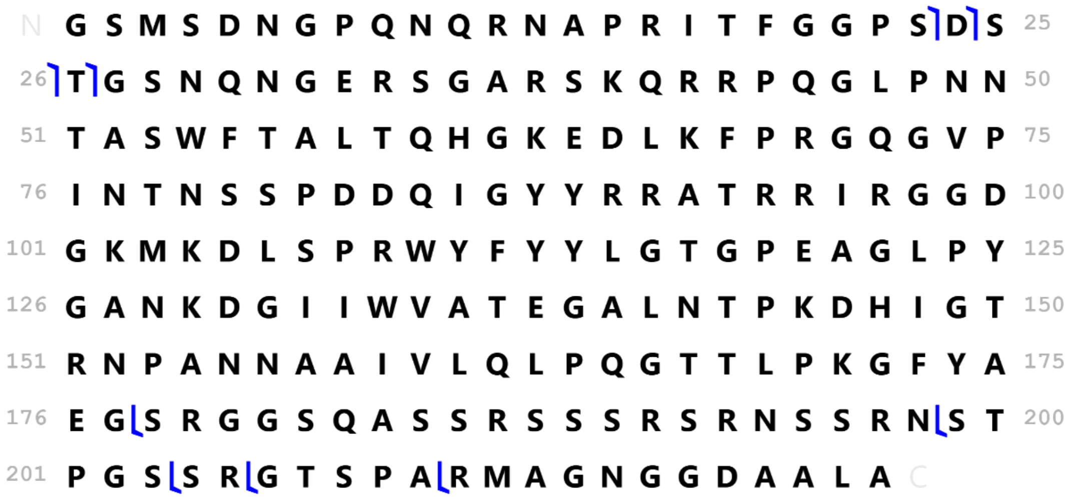

**Deconvoluted mass of charge state distribution**: 29,402 ± 0.6*
**Tentative Assignment**: $N_{1\text{-}273}$

| Candidate Sequence | Sequence Mass (Da) | Precursor Charge State | Fragments matched | P-score | Sequence Coverage (%) |
|---|---|---|---|---|---|
| $N_{FL}$ | 45,769.83 | | | | |
| | | 10+ | 0 | 1 | 0 |
| | | 11+ | 3 | 2.0e-2 | 0 |
| $N_{1\text{-}272}$ | 29,311.35 | | | | |
| | | 10+ | 0 | 1 | 0 |
| | | 11+ | 3 | 2.0e-2 | 0 |
| **$N_{1\text{-}273}$** | 29,382.43 | | | | |
| | | 10+ | 0 | 1 | 0 |
| | | **11+** | 6 | 2.90e-5 | 2 |
| $N_{1\text{-}274}$ | 29,529.60 | | | | |
| | | 10+ | 0 | 1 | 0 |
| | | 11+ | 3 | 2.0e-2 | 0 |
| $N_{148\text{-}419}$ | 29,433.89 | | | | |
| | | 10+ | 1 | 0.4 | 0 |
| | | 11+ | 0 | 1 | 0 |
| $N_{149\text{-}419}$ | 29,277.70 | | | | |
| | | 10+ | 2 | 0.095 | 1 |
| | | 11+ | 0 | 1 | 0 |

N G S M S D N G P Q N Q R N A P R I T F G G P S D S 25
26 T G S N Q N G E R S G A R S K Q R R P Q G L P N N 50
51 T A S W F T A L T Q H G K E D L K F P R G Q G V P 75
76 I N T N S S P D D Q I G Y Y R R A T R R I R G G D 100
101 G K M K D L S P R W Y F Y Y L G T G P E A G L P Y 125
126 G A N K D G I I W V A T E G A L N T P K D H I G T 150
151 R N P A N N A A I V L Q L P Q G T T L P K G F Y A 175
176 E G S R G G S Q A S S R S S S R S R N S S R N S T 200
201 P G S S R G T S P A R M A G N G G D A A L A L L L 225
226 L D R L N Q L E S K M S G K G Q Q Q Q G Q T V T K 250
251 K S A A E A S K K P R Q K R T A T K A Y N V T Q A C

*The 20 Da difference in deconvoluted mass and sequence mass may arise from the oxidation of methionine residues and/or unaccounted metal ion cofactors.

**Deconvoluted mass of charge state distribution**: 28,735 ± 2*
**Tentative Assignment**: $N_{156\text{-}419}$

| Candidate Sequence | Sequence Mass (Da) | Precursor Charge State | Fragments matched | P-score | Sequence Coverage (%) |
|---|---|---|---|---|---|
| $N_{FL}$ | 45,769.83 | | | | |
| | | 10+ | 1 | 0.54 | 0 |
| | | 11+ | 0 | 1 | 0 |
| $N_{1\text{-}266}$ | 28,634.62 | | | | |
| | | 10+ | 0 | 1 | 0 |
| | | 11+ | 0 | 1 | 0 |
| $N_{1\text{-}267}$ | 28,705.70 | | | | |
| | | 10+ | 0 | 1 | 0 |
| | | 11+ | 0 | 1 | 0 |
| $N_{1\text{-}268}$ | 28,868.88 | | | | |
| | | 10+ | 0 | 1 | 0 |
| | | 11+ | 0 | 1 | 0 |
| $N_{154\text{-}419}$ | 28,881.30 | | | | |
| | | 10+ | 1 | 0.54 | 0 |
| | | 11+ | 0 | 1 | 0 |
| $N_{155\text{-}419}$ | 28,767.20 | | | | |
| | | 10+ | 1 | 0.54 | 0 |
| | | 11+ | 0 | 1 | 0 |
| **$N_{156\text{-}419}$** | **28,696.12** | | | | |
| | | **10+** | **2** | **0.18** | **1** |
| | | 11+ | 0 | 1 | 0 |

N A I V L Q L P Q G T T L P K G F Y A E G S R G G S 25
26 Q A S S R S S S R S R N S S R N S T P G S S R G T 50
51 S P A R M A G N G G D A A L A L L L L D R L N Q L 75
76 E S K M S G K G Q Q Q Q G Q T V T K K S A A E A S 100
101 K K P R Q K R T A T K A Y N V T Q A F G R R G P E 125
126 Q T Q G N F G D Q E L I R Q G T D Y K H W P Q I A 150
151 Q F A P S A S A F F G M S R I G M E V T P S G T W 175
176 L T Y T G A I K L D D K D P N F K D Q V I L L N K 200
201 H I D A Y K T F P P T E P K K D K K K K A D E T Q 225
226 A L P Q R Q K K Q Q T V T L L P A A D L D D F S K 250
251 Q L Q Q S M S S A D S T Q A C

*The 40 Da difference in deconvoluted mass and sequence mass may arise from the oxidation of methionine residues and/or unaccounted metal ion cofactors. Limited peptide coverage was achieved by top-down MS due to low initial ion intensity.

**Deconvoluted mass of charge state distribution**: 42,922 ± 1

**Tentative Assignment**: $N_{1-392}$

| Candidate Sequence | Sequence Mass (Da) | Precursor Charge State | Fragments matched | P-score | Sequence Coverage (%) |
|---|---|---|---|---|---|
| $N_{FL}$ | 45,769.83 | 13+ | 0 | 1 | 0 |
| | | 12+ | 1 | 1.8e-1 | 0 |
| $N_{1-391}$ | 42,718.50 | 13+ | 0 | 1 | 0 |
| | | 12+ | 1 | 1.8e-1 | 0 |
| $N_{1-392}$ | 42,918.74 | 13+ | 12 | 6.1e-14 | 4 |
| | | 12+ | 10 | 2.1e-11 | 3 |
| $N_{1-393}$ | 43,019.84 | 13+ | 1 | 0.19 | 0 |
| | | 12+ | 2 | 1.8e-2 | 1 |
| $N_{24-419}$ | 43,096.02 | 13+ | 0 | 1 | 0 |
| | | 12+ | 0 | 1 | 0 |

N G S M S D N G P Q N Q R N A P R I T F G G P S D S 25
26 T G S N Q N G E R S G A R S K Q R R P Q G L P N N 50
51 T A S W F T A L T Q H G K E D L K F P R G Q G V P 75
76 I N T N S S P D D Q I G Y Y R R A T R R I R G G D 100
101 G K M K D L S P R W Y F Y Y L G T G P E A G L P Y 125
126 G A N K D G I I W V A T E G A L N T P K D H I G T 150
151 R N P A N N A A I V L Q L P Q G T T L P K G F Y A 175
176 E G S R G G S Q A S S R S S S R S R N S S R N S T 200
201 P G S S R G T S P A R M A G N G G D A A L A L L L 225
226 L D R L N Q L E S K M S G K G Q Q Q Q G Q T V T K 250
251 K S A A E A S K K P R Q K R T A T K A Y N V T Q A C

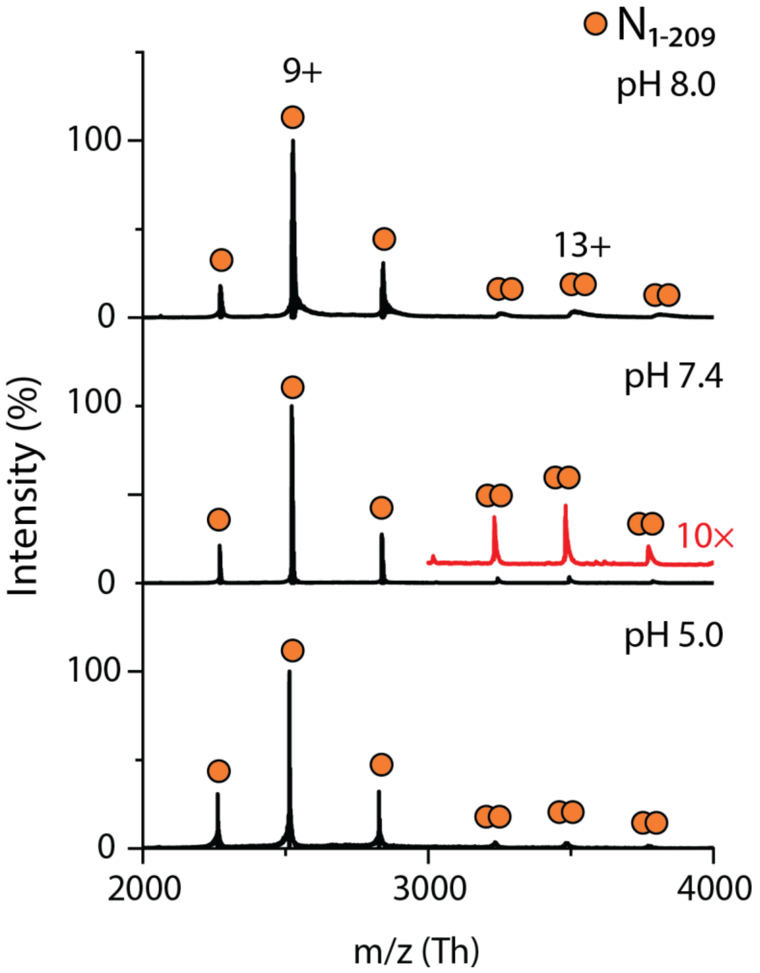


**Figure S3.** The oligomeric state of $N_{1\text{-}209}$ is not influenced by pH. Mass spectra of $N_{1\text{-}209}$ measured at pH 8.0, 7.4 and 5.0 (top to bottom). Two charge state distributions are observed that correspond to monomers and dimers of $N_{1\text{-}209}$, centered at 9+ and 13+, respectively. The mass spectrum recorded at pH 7.4 was magnified 10× at *m/z* 3000 and offset for clarity (red trace) to highlight the low abundance distribution of $N_{1\text{-}209}$ dimer. The deconvoluted masses of each distribution can be found in Table S2.

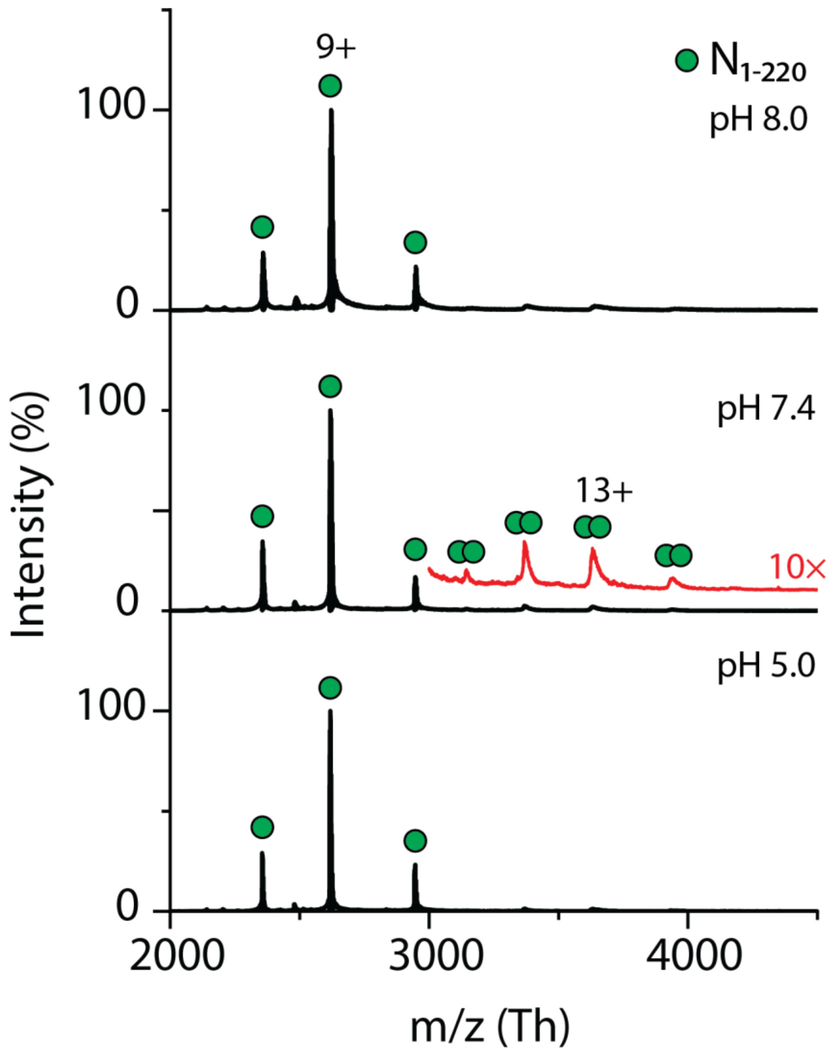


**Figure S4.** The oligomeric state of $N_{1\text{-}220}$ is not influenced by pH. Mass spectra of $N_{1\text{-}220}$ measured at pH 8.0, 7.4 and 5.0 (top to bottom). Two charge state distributions are observed that correspond to monomers and dimers of $N_{1\text{-}220}$, centered at 9+ and 13+, respectively. The mass spectrum recorded at pH 7.4 was magnified 10× at *m/z* 3000 and offset for clarity (red trace) to highlight the low abundance distribution of $N_{1\text{-}220}$ dimer. The deconvoluted masses of each distribution can be found in Table S2.

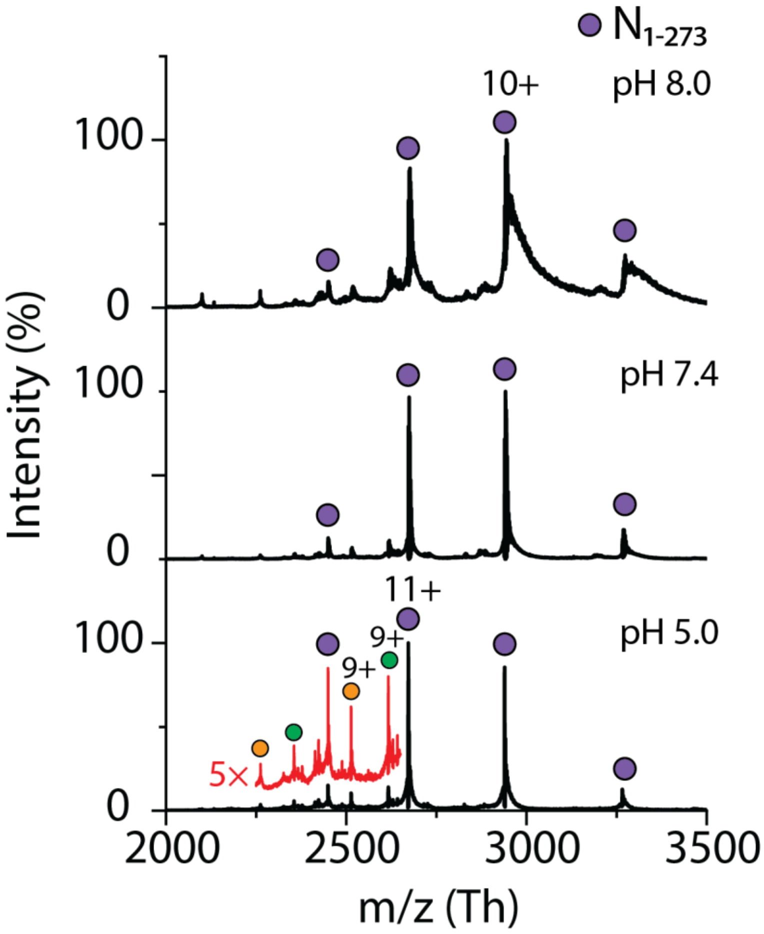


**Figure S5.** The oligomeric state of $N_{1\text{-}273}$ is not influenced by pH. Mass spectra of $N_{1\text{-}273}$ measured at pH 8.0, 7.4 and 5.0 (top to bottom). One charge state distribution is observed that correspond to monomeric $N_{1\text{-}273}$, centered at 10+ at pH 8.0 and 7.4, and 11+ at pH 5.0. The mass spectrum recorded at pH 5.0 was magnified 5× at *m/z* 2250-2650 and offset for clarity (red trace) to highlight the low abundance charge state distributions corresponding to $N_{1\text{-}209}$ and $N_{1\text{-}220}$ proteoforms resulting from the continued proteolytic cleavage of $N_{1\text{-}273}$. The deconvoluted masses of each distribution can be found in Table S2.

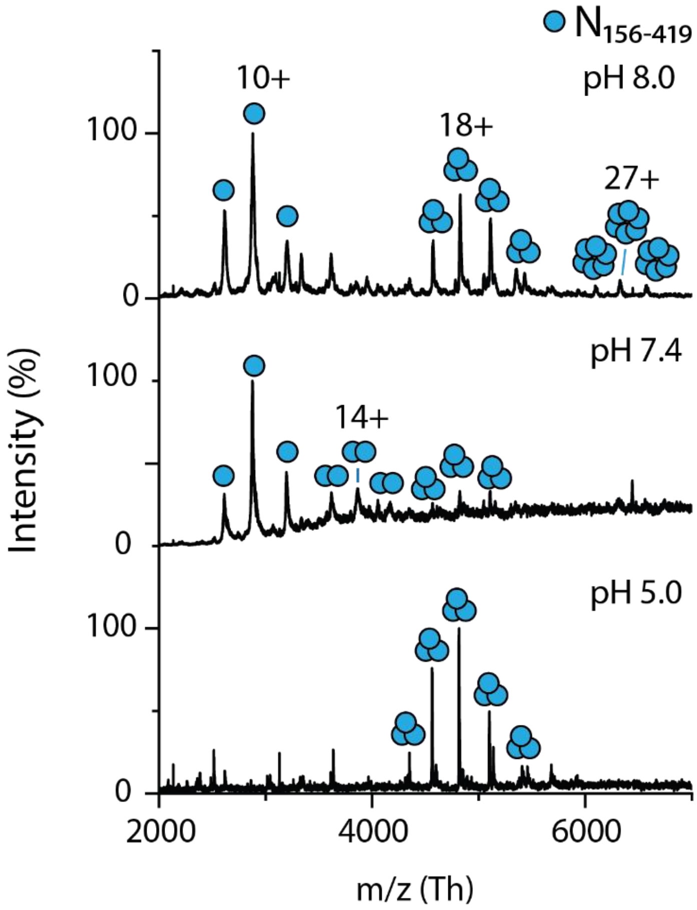


**Figure S6.** pH influences the oligomeric state of $N_{156\text{-}419}$. Mass spectra of $N_{156\text{-}419}$ measured at pH 8.0, 7.4, and 5.0 (top to bottom). $N_{156\text{-}419}$ appears to exhibit a pH-dependence on oligomeric state. At pH 8.0, the mass spectrum reveals three charge state distributions corresponding to monomers, trimers, and hexamers centered charge states 10+, 18+ and 27+, respectively. At pH 7.4, three charge state distributions corresponding to monomers, dimers and trimers are observed. At pH 5.0, only one charge state distribution is observed, indicating that $N_{156\text{-}419}$ is exclusively trimeric at pH 5.0. The deconvoluted masses of each distribution can be found in Table S2.

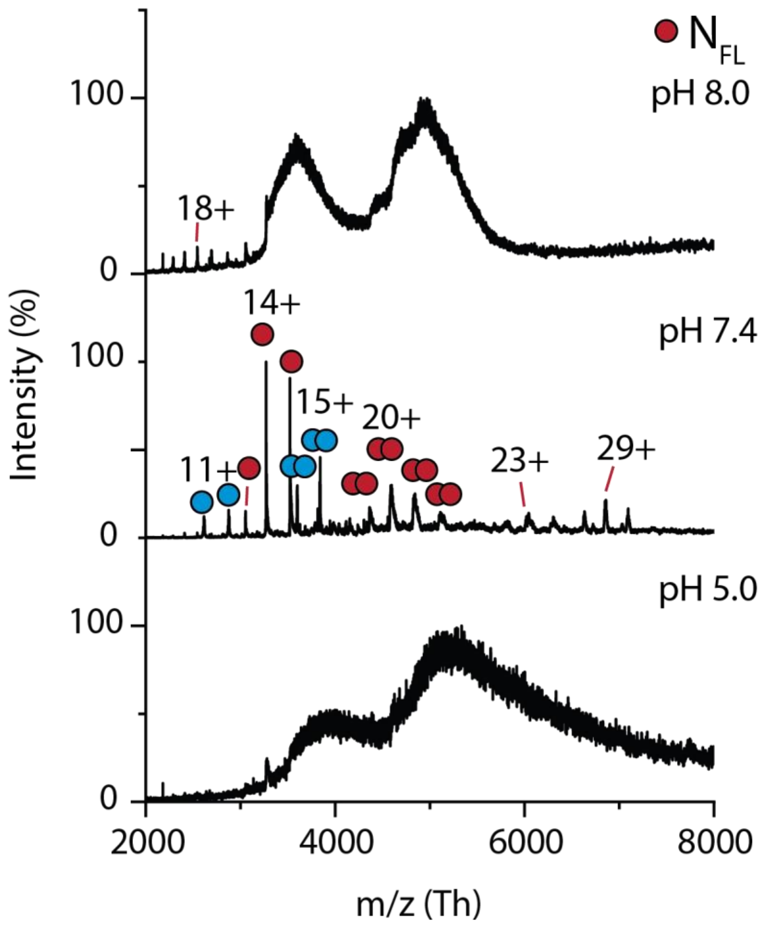


**Figure S7**. Effect of pH on the oligomeric state of N$_{FL}$. Mass spectra of N$_{FL}$ measured at pH 8.0, 7.4 and 5.0 (top to bottom). At pH 7.4, several charge state distributions are observed that correspond to monomers and dimers of NFL (red circles with charges 14+ and 20+, respectively), monomers and dimers of N156-419 (blue circles with charges of 11+ and 15+, respectively) and two remaining charge state distributions centered at 23+ and 29+ that may correspond to hetero-oligomers of N$_{FL}$ and N proteoforms. The deconvoluted masses of each distribution can be found in Table S2. The mass spectra at pH 5.0 and 8.0 reveal broadened and featureless peaks suggesting that it is likely aggregated. A low abundance series of highly charged peaks centered at 18+ at pH 8.0 indicates some protein unfolding.

**Table S2**. Deconvoluted and expected masses of assigned charge state distributions in Figures S4-S7

| Protein | Deconvoluted Mass[a] ± s.d. (Da) | Sequence Mass (Da) |
|---|---|---|
| $N_{1-209}$ | 22,611 ± 1 | 22,612.71 |
| $N_{1-209}$ dimer | 45,225 ± 2 | 45,225.42 |
| $N_{1-220}$ | 23,540 ± 0.3 | 23,541.73 |
| $N_{1-220}$ dimer | 47,106 ± 1 | 47,083.46 |
| $N_{1-273}$ | 29,402 ± 0.6 | 29,382.43 |
| $N_{156-419}$ | 28,735 ± 2 | 28,696.12 |
| $N_{156-419}$ dimer | 57,534 ± 1 | 57,392.24 |
| $N_{156-419}$ trimer | 86,700 ± 13 | 86,088.36 |
| $N_{156-419}$ hexamer | 170,813 ± 34 | 172,176.72 |
| $N_{FL}$ | 45,769 ± 2 | 45,769.83 |
| $N_{FL}$ dimer | 91,537 ± 2 | 91,539.66 |
| Distribution centered at +23 | 144,958 ± 28 | -- |
| Distribution centered at +29 | 205,634 ± 23 | -- |

[a] Determined using three adjacent charge states

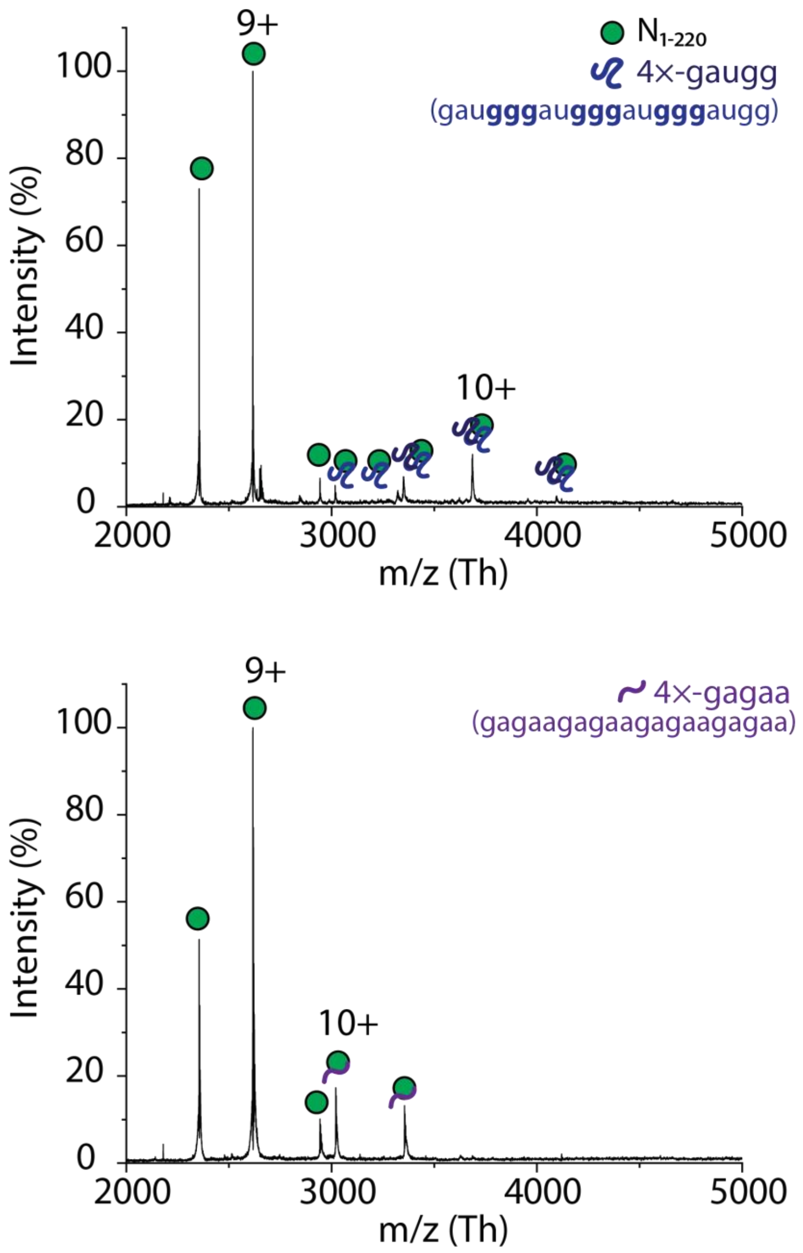


**Figure S8**. The RNA sequence influences binding stoichiometry to $N_{1\text{-}220}$. (A) Mass spectrum of $N_{1\text{-}209}$ after incubation with 4×-GAUGG RNA oligonucleotides in a molar ratio of 1:4. The most abundant charge state distribution corresponds to monomeric $N_{1\text{-}220}$ centered at the 9+ charge state. Two additional charge state distributions are observed above *m/z* 3000 that correspond to one and two RNA oligonucleotides bound to $N_{1\text{-}220}$. (B) Mass spectrum of $N_{1\text{-}220}$ after incubation with 4×-GAGAA RNA oligonucleotides in a molar ratio of 1:4. The most abundant charge state distribution corresponds to monomeric $N_{1\text{-}220}$ centered at the 9+ charge state. An additional, low abundant charge state series is observed between *m/z* 3000 and 3500 that corresponds to one RNA oligonucleotide bound to $N_{1\text{-}220}$.

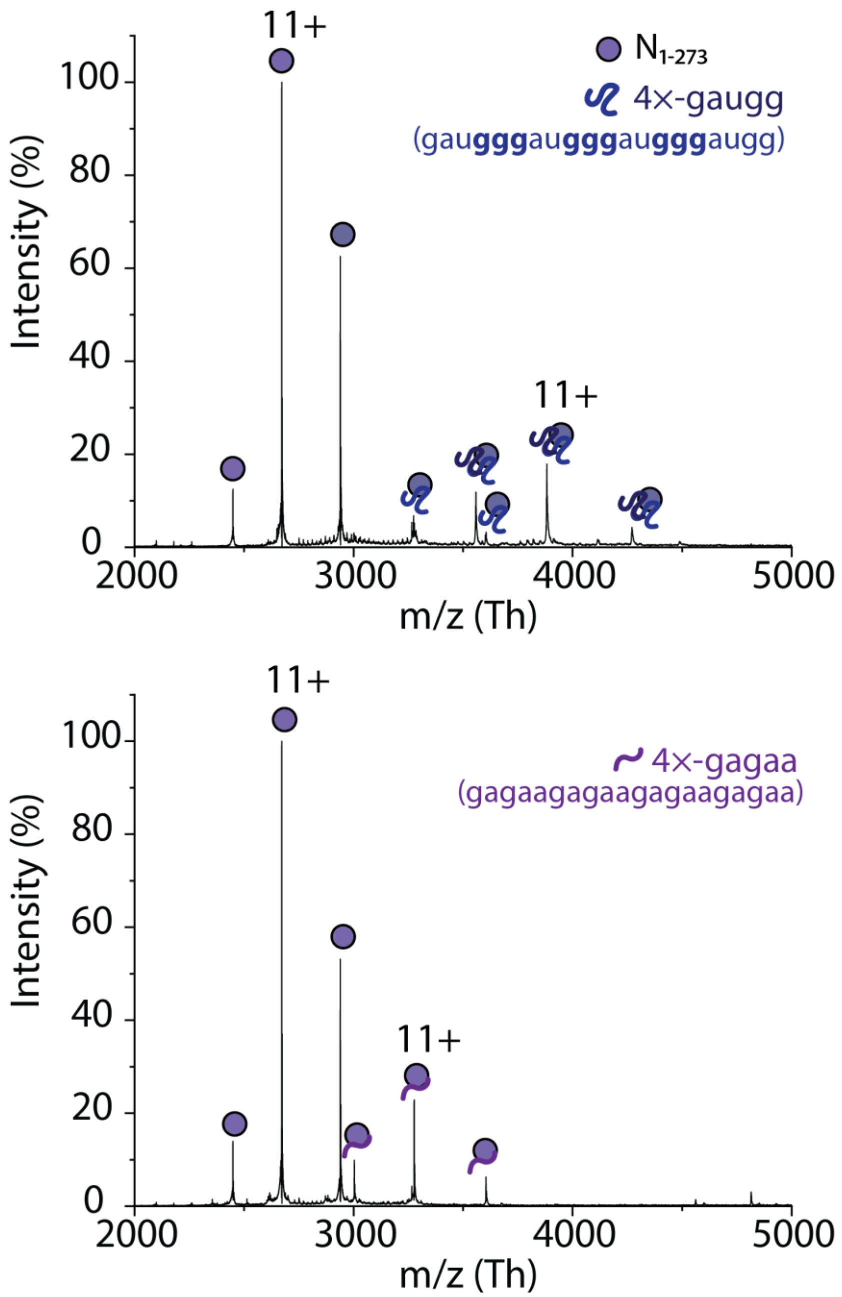


**Figure S9**. The RNA sequence influences binding stoichiometry to $N_{1\text{-}273}$. (A) Mass spectrum of $N_{1\text{-}273}$ after incubation with 4×-GAUGG RNA oligonucleotides in a molar ratio of 1:4. The most abundant charge state distribution corresponds to monomeric $N_{1\text{-}273}$ centered at the 11+ charge state. Two additional charge state distributions are observed above *m/z* 3000 that correspond to one and two RNA oligonucleotides bound to $N_{1\text{-}273}$. (B) Mass spectrum of $N_{1\text{-}273}$ after incubation with 4×-GAGAA RNA oligonucleotides in a molar ratio of 1:4. The most abundant charge state distribution corresponds to monomeric $N_{1\text{-}273}$ centered at the 11+ charge state. An additional, low abundant charge state series is observed between *m/z* 3000 and 4000 and centered at a charge state of 11+ that corresponds to one RNA oligonucleotide bound to $N_{1\text{-}273}$.

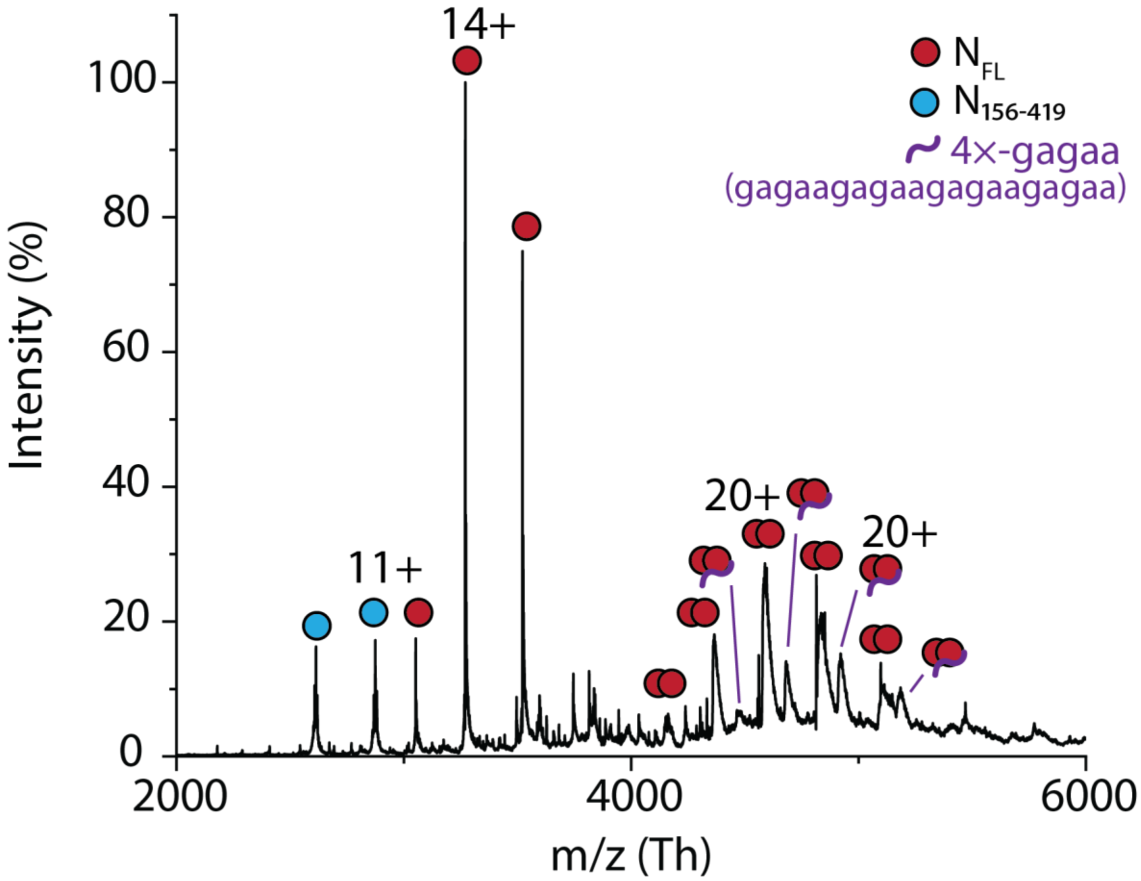


**Figure S10**. Mass spectrum of $N_{FL}$ after incubation with 4×-GAGAA RNA oligonucleotides in a molar ratio of 1:4. Monomers and dimers of $N_{FL}$ (red circles) and $N_{156-419}$ (blue circles) are observed. An additional peak series between 4300 and 5500 *m/z* corresponds to one 4×-gagaa RNA oligonucleotide bound to $N_{FL}$ dimer.

**Table S3**. Deconvoluted and sequence masses of N protein-RNA complexes.

| Protein/Complex | Deconvoluted Mass ± s.d. (Da) | Sequence Mass (Da) |
|---|---|---|
| 4×-GAUGG RNA | -- | 6,622.00 |
| 4×-GAGAA RNA | -- | 6,650.20 |
| $N_{1-220}$ + one 4×-GAUGG RNA | 30,162 ± 1.0 | 30,163.73 |
| $N_{1-220}$ + two 4×-GAUGG RNA | 36,847 ± 13 | 36,785.73 |
| $N_{1-220}$ + one 4×-GAGAA RNA | 30,191 ± 0.2 | 30,191.93 |
| $N_{1-273}$ + one 4×-GAUGG RNA | 36,022 ± 27 | 36,004.43 |
| $N_{1-273}$ + two 4×-GAUGG RNA | 42,696 ± 1.0 | 42,626.43 |
| $N_{1-273}$ + one 4×-GAGAA RNA | 36,031 ± 0.5 | 36,032.63 |
| $N_{FL}$ dimer + 4×-GAGAA RNA | 98,255 ± 46 | 98,189.86 |

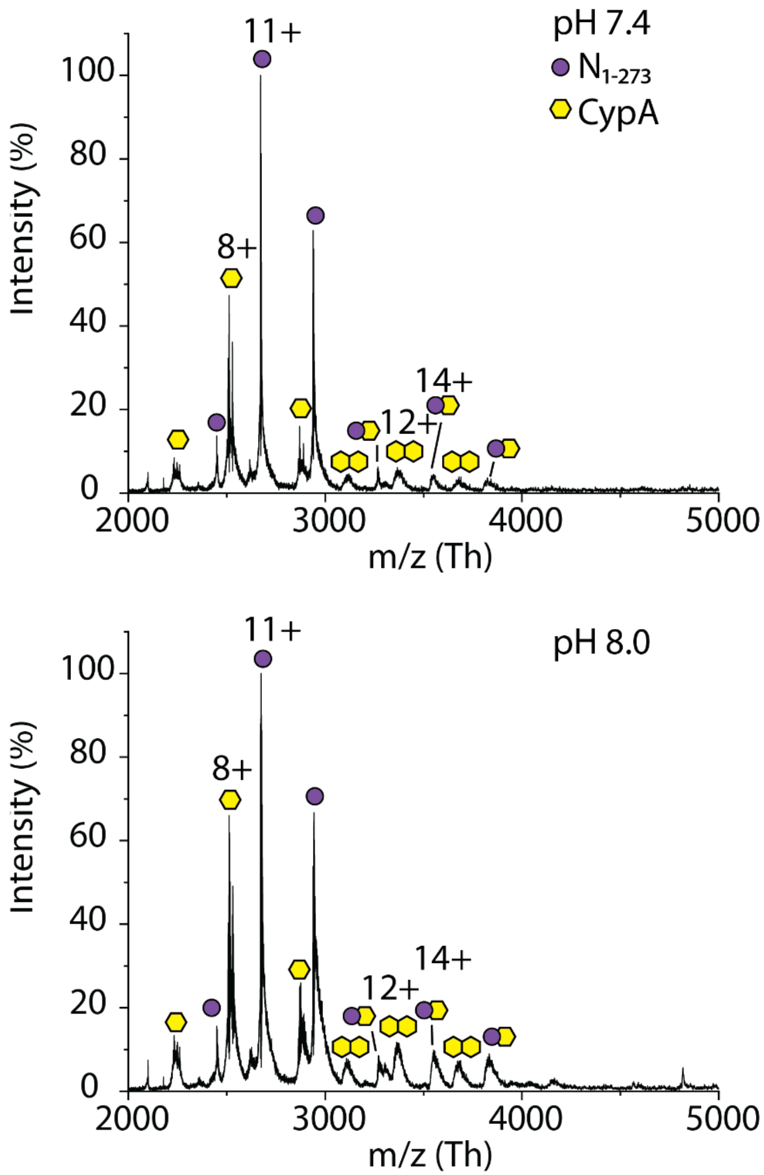


**Figure S11**. $N_{1-273}$ directly interacts with cyclophilin A. Mass spectra show $N_{1-273}$ after incubation with cyclophilin A (CypA) in a 1:1 molar ratio at pH 7.4 (top) and pH 8.0 (bottom). In both spectra, four distinct charge state distributions are observed: (i) a high abundant charge state distribution centered at 11+ that corresponds to N1-273 monomer, (ii) a second abundant charge state distribution centered at 8+ that corresponds to CypA, (iii) a low abundant distribution from *m/z* 3000-4000 centered at 12+ that corresponds to CypA dimers, and (iv) a second low abundant distribution between *m/z* 3000 and 4000 centered at 14+ that corresponds to $N_{1-273}$ bound to cypA.

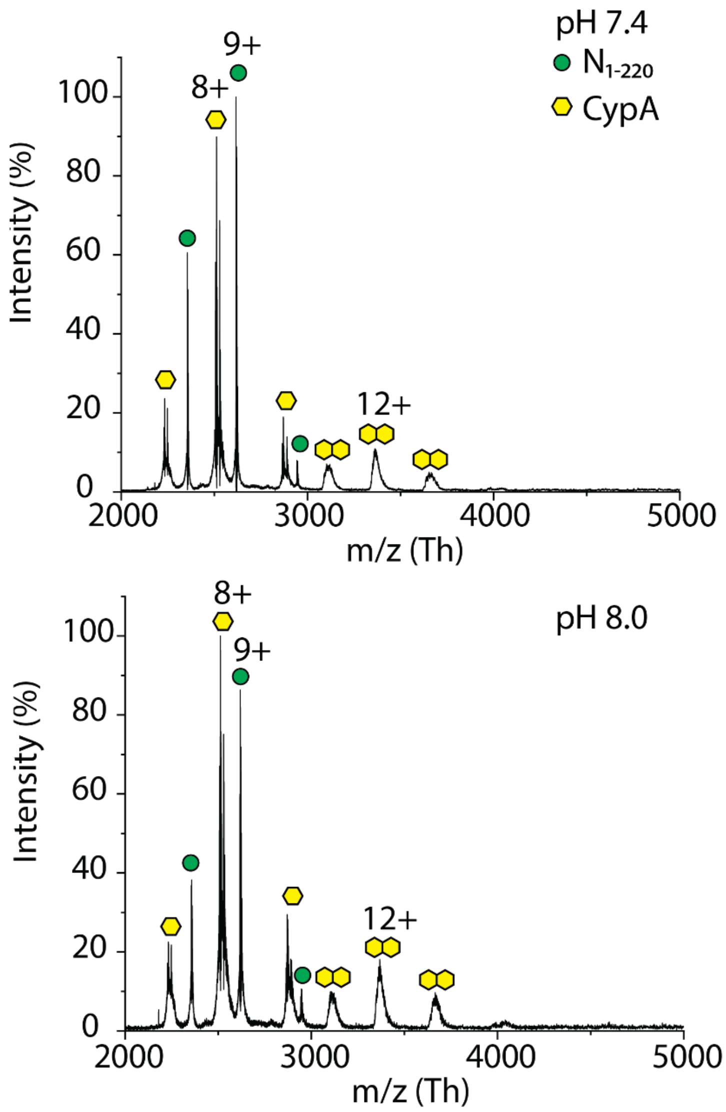


**Figure S12**. $N_{1\text{-}220}$ does not interact with cyclophilin A. Mass spectra show $N_{1\text{-}220}$ after incubation with cyclophilin A (CypA) in a 1:1 molar ratio at pH 7.4 (top) and pH 8.0 (bottom). In both spectra, three charge state distributions are observed: (i) a high abundant charge state distribution centered at 9+ that corresponds to $N_{1\text{-}220}$ monomers, (ii) a second highly abundant charge state distribution centered at 8+ that corresponds to CypA, and (iii) a low abundant charge state distribution at *m/z* >3000 and centered at 12+ that corresponds to CypA dimers. No interaction is observed between $N_{1\text{-}220}$ and CypA.

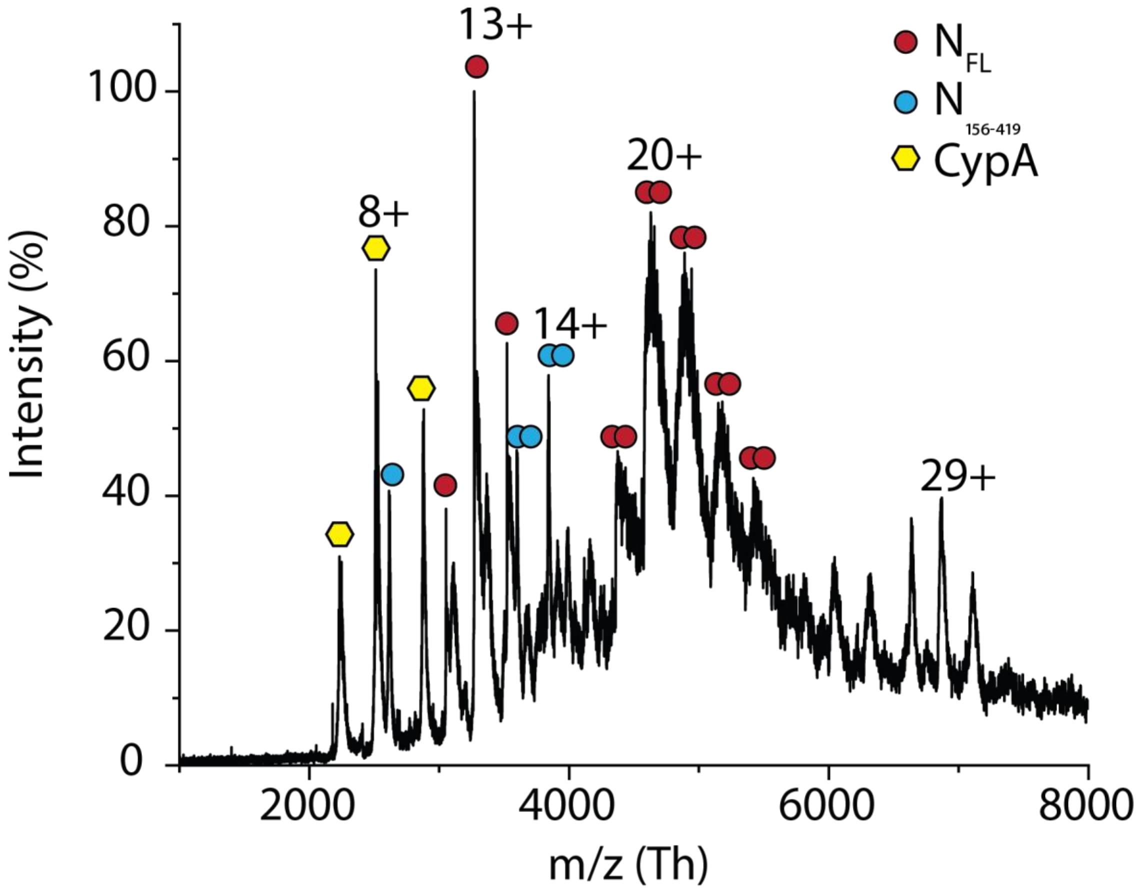


**Figure S13**. $N_{FL}$ does not interact with cyclophilin A. Mass spectrum shows $N_{FL}$ after incubation with cyclophilin A (CypA) in a 1:1 molar ratio. Charge state distributions are observed for monomeric CypA centered at a charge state of 8+ (yellow hexagons), NFL monomers and dimers centered at 13+ and 20+, respectively (red circles), $N_{156\text{-}419}$ monomers and dimers (blue circles), and a charge state distribution at *m/z* >6000 centered at 29+. No obvious interaction between CypA and $N_{FL}$ is observed.

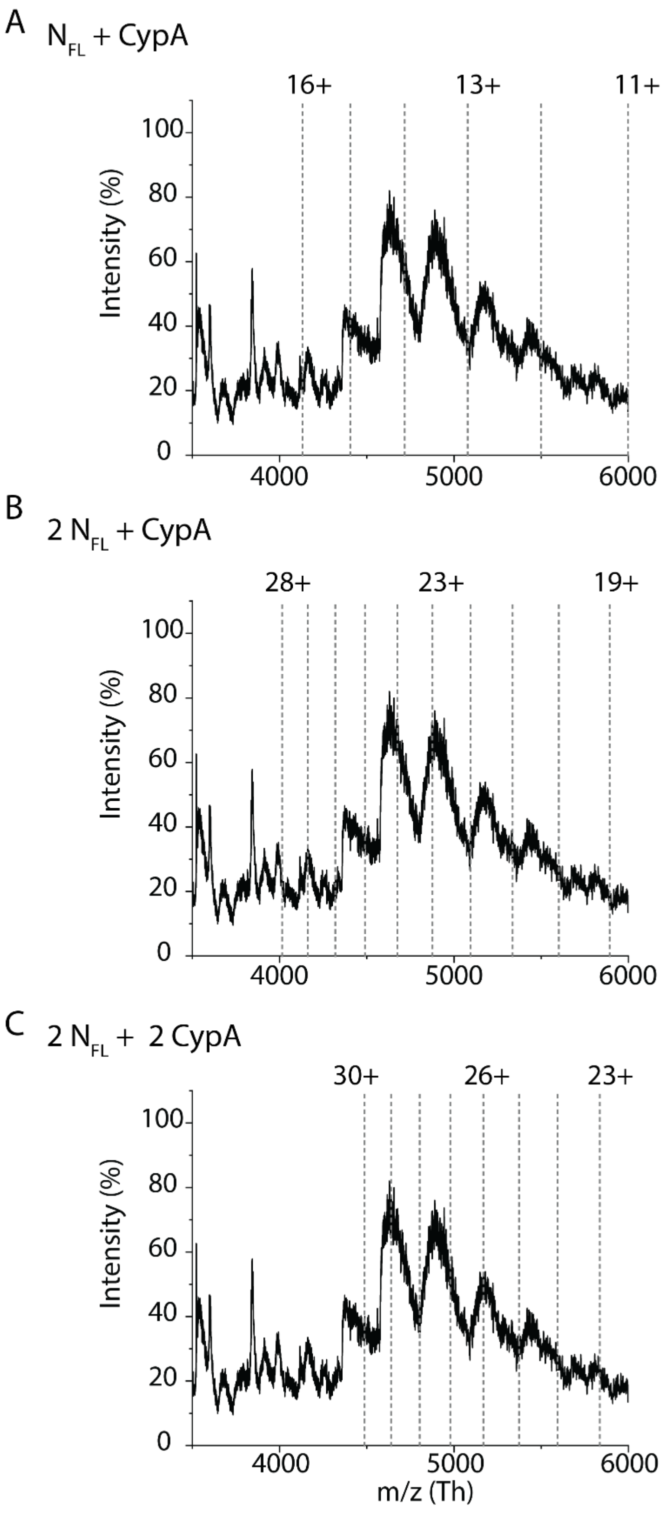


**Figure S14**. Theoretical peak series for $N_{FL}$-CypA complex formation overlaid onto the measured mass spectrum for $N_{FL}$ + CypA. The grey dashed lines represent the expected *m/z* values for (A) a complex of $N_{FL}$ bound to CypA, (B) $N_{FL}$ dimers bound to CypA, and (C) $N_{FL}$ dimer bound to CypA dimers. No observed peaks match the anticipated distributions, therefore $N_{FL}$ and CypA do not strongly interact, if at all.

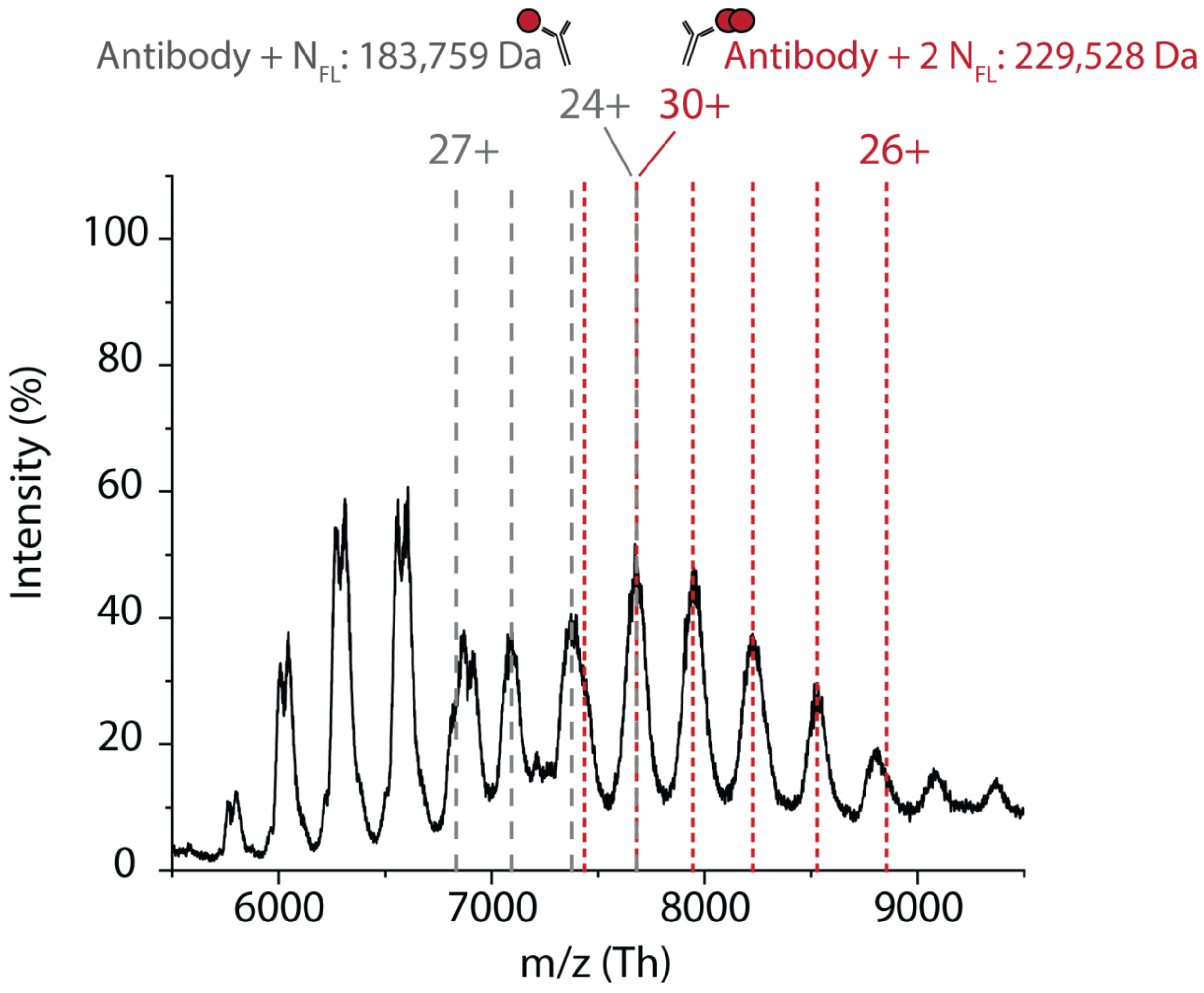


**Figure S15**. Calculated charge state distributions for one and two $N_{FL}$ bound to a monoclonal antibody. The grey dashed lines represent the *m/z* values for charge states 24+ through 27+ for the antibody bound to one $N_{FL}$ corresponding to a mass of 183,759 Da. The red dashed lines represent the *m/z* values for charge states 26+ through 30+ for the antibody bound to two $N_{FL}$ corresponding to a mass of 229,528 Da. The expected masses were calculated using the lowest deconvoluted mass of the antibody (137,990 Da). The deconvoluted masses for the measured distributions (black trace) are 183,931 ± 102 Da and 229,984 ± 36 Da, respectively.
